# Developmentally arrested precursors of pontine neurons establish an embryonic blueprint of the *Drosophila* central complex

**DOI:** 10.1101/291831

**Authors:** Ingrid V. Andrade, Nadia Riebli, Bao-Chau M. Nguyen, Jaison J. Omoto, Richard D. Fetter, Albert Cardona, Volker Hartenstein

## Abstract

Serial electron microscopic analysis shows that the *Drosophila* brain at hatching possesses a large fraction of developmentally arrested neurons with a small soma, heterochromatin-rich nucleus, and unbranched axon lacking synapses. We digitally reconstructed all 812 “ small undifferentiated” (SU) neurons on both hemispheres and assigned them to the known brain lineages. 54 SU neurons belonging to the DM1-4 lineages, which generate all columnar neurons of the central complex, form an embryonic nucleus of the fan-shaped body (FB). These “ FB pioneers” develop into a specific class of bi-columnar elements, the pontine neurons. Even though later born, unicolumnar DM1-4 neurons fasciculate with the FB pioneers, selective ablation of these cells did not result in gross abnormalities of the trajectories of unicolumnar neurons, indicating that axonal pathfinding of the two systems is controlled independently. Our comprehensive spatial and developmental analysis of the SU neuron adds to our understanding of the establishment of neuronal circuitry.

## Introduction

The central complex (CX) of the insect brain plays an important role in a variety of different behaviors, including the 1ne control of motor movement and spatial orientation (Martin et al., 1999; Neuser et al., 2008; Bender et al., 2010; Kahsai et al., 2010; Ofstad et al. 2011; Pfeiffer et al., 2014). Recent studies indicate that the CX harbors dynamic neural activity that integrates the animal’ s external and internal environment (Seelig and Jayaraman, 2015; Green et al., 2017; Turner-Evans et al., 2017). Anatomical studies have started to reveal the neuronal connectivity that underlies CX function. The CX is comprised of four major compartments, including (from anterior to posterior) the ellipsoid body (EB), fan-shaped body (FB) with noduli (NO), and protocerebral bridge (PB; Strausfeld, 1976; Hanesch et al., 1989; Fig.1). CX circuitry is dominated by an orthogonal scaffold of transversally oriented (“tangential”) widefield neurons, and longitudinally oriented columnar small field neurons. Tangential neurons, whose fibres are directed parallel to the length axis of the CX neuropils (a, b, c in Fig.1), provide input to the CX from other brain areas. Best understood among these input neurons are the TL-neurons in locust (el Jundi et al., 2014) and their *Drosophila* counterparts, the R-neurons, that conduct retinotopically ordered visual information to the ellipsoid body (Seelig and Jayaraman, 2013; Fig.1). Columnar neurons, which interconnect the different CX neuropils along the antero-posterior axis, are characterized by highly localized dendritic and axonal endings in narrow volumes (“ columns”) of the respective compartments. Most classes of columnar neurons, within a given CX neuropil, are confined to a single column (“unicolumnar neurons”; Fig.1). Projections are characterized by a strict homotopic order, whereby columns within the lateral half of the PB are connected to columns of the ipsilateral PB and EB, and medial PB columns project to the contralateral FB and EB (Fig.1). One class of columnar neurons, the so-called pontine neurons of the fan-shaped body (Hanesch et al., 1989; Young and Armstrong, 2010), behave differently. Their projection is restricted to the FB, interconnecting two FB columns located on either side of the midline (“bicolumnar neurons”; Fig.1).

**Figure 1.**
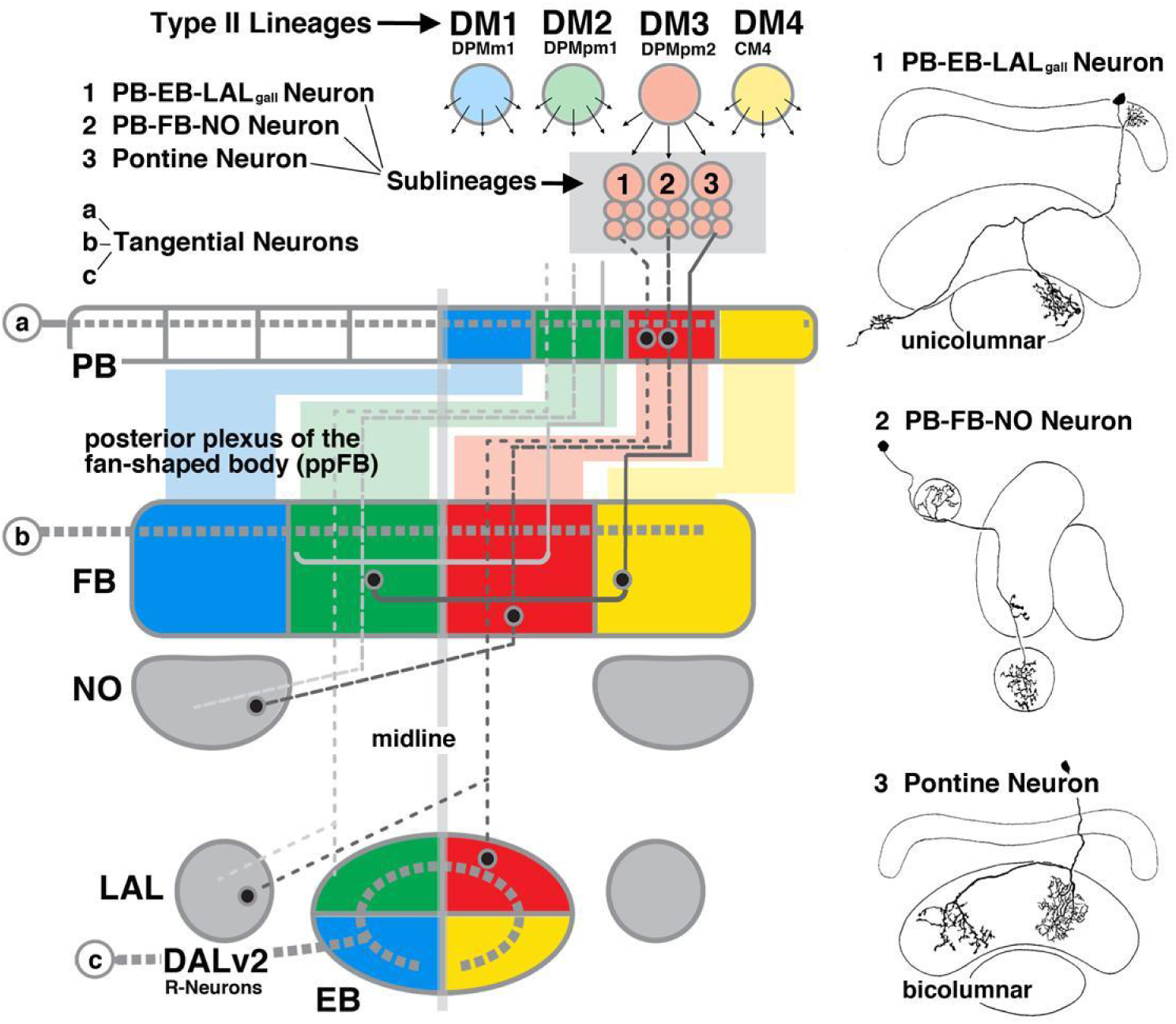
Connectivity and cell types of the *Drosophila* central complex in the context of neural lineages. Left: Schematic of the central complex, showing its main compartments (PB: protocerebral bridge; FB: fan-shaped body; NO: noduli; EB: ellipsoid body; LAL: lateral accessory lobe). Four bilateral pairs of lineages, DM1-DM4, located in the posterior brain cortex (top), generate the columnar neurons whose axons (grey lines oriented along the vertical axis; 1lled circles symbolize terminal arborizations) interconnect the compartments of the central complex along the antero-posterior axis in a strict homotopic order. The spatial pattern of DM1-4 is re2ected in the position at which their corresponding tracts enter and terminate within the CX, as indicated by the coloring scheme. The four lineages subdivide the CX neuropils into four quadrants; projections of all neurons belonging to one lineage (with the exception of the pontine neurons) are con1ned to one quadrant. The lineage located furthest laterally, DM4 (yellow), innervates the ipsilateral lateral quadrant; DM3 projects ipsilateral medially (red), DM2 contralateral medially (green), and DM1 contralateral laterally (blue). Columnar neurons are further subdivided into numerous classes according to their projection. The three examples shown include the PB-EB-LALgall neurons (1; hatched line with wide gaps), PB-FB-NO neurons (2; hatched line with narrow gaps) and pontine neurons (3; solid line). These classes may encompass different sublineages, as indicated by small arrows radiating out from the neuroblasts (top). Pontine neurons, as the only class of DM1-4-derived neurons, are restricted to the FB where they interconnect two quadrants. Apart from the columnar neurons, tangential neurons (a, b, c; thick hatched horizontal lines) project across all columns of the CX compartments, and provide input from the lateral brain. Right: Line drawings of the central complex in dorsal view (top, bottom) or sagittal view (center), depicting examples of two unicolumnar neurons (1:PB-EB-LALgall neuron; 2: PB-FB-NO neuron) and a bicolumnar pontine neuron (3; from Hanesch et al., 1989, with permission).

Developmental studies provide a valuable approach to unravel the circuitry of the brain, including the central complex. A hallmark of the *Drosophila* brain is its composition of invariant neuronal and glial lineages, originating from stem cells (neuroblasts) that appear in the early embryo. Embryonic neuroblasts express speci1c combinations of transcription factors (TFs), which are thought to provide each lineage with the information needed to shape the connectivity of its neurons (Urbach and Technau, 2004; Brody and Odenwald, 2005; Kohwi and Doe, 2014). As a result, lineages become structural modules: Neurons of the same lineage generally project together in one or two 1ber tracts, and form synapses in specific, spatially restricted brain compartments. Neuron classes of the CX conform well to the lineage principle. For example, the R-neurons of the ellipsoid body are derived from one lineage, DALv2 (also called EBa1) (Larsen et al., 2009; Spindler and Hartenstein, 2011; Wong et al., 2013; Yang et al., 2013; Yu et al., 2013; Omoto et al., 2017; 2018). Sublineages of DALv2 born at different times further tile the bulb (BU) and EB into discrete layers (Omoto et al., 2018). The columnar neurons of the CX are produced by four pairs of lineages located in the dorsomedial brain, called DM1/DPMm1, DM2/DPMpm1, DM3/DPMpm2, DM4/CM4 (called DM1-DM4 henceforward; Ito and Awasaki, 2008; Bello et al., 2008; Ito et al., 2013; Wong et al., 2013; Yang et al., 2013; Yu et al., 2013). The spatial pattern of these lineages is reflected in the position at which their corresponding tracts enter and terminate within the CX. In this manner, the four lineages subdivide the CX neuropils into four evenly sized quadrants, as illustrated in Fig.1.

The brain of *Drosophila* and other holometabolous insects arises in two distinct phases. During the 1rst phase, neuroblasts of the embryo produce a relatively small set of primary neurons which differentiate and form the larval brain. Most neuroblasts then enter a dormant phase that lasts towards the end of the first larval instar. Subsequently they reactivate and produce secondary, adult speci1c neurons. These cells form axon bundles that form a “blue print” of connections later established within the adult brain. Differentiation is delayed until metamorphosis, when secondary neurons, along with re-modeled primary neurons, extend axonal and dendritic branches and form synapses. In general, primary and secondary neurons of a given lineage show fundamental structural similarities, whereby projections of secondary neurons follow those of earlier formed primary neurons (Larsen et al., 2009). Remarkably, the central complex, as de1ned anatomically for the adult, is the one major set of compartments of the 2y brain that lacks an obvious counterpart in the larva. Thus, all tangential and columnar neurons with their highly ordered connections outlined above are secondary neurons born in the larva, prompting the question of what guidance mechanisms control CX connectivity, and what part primary neurons play during this process.

Previous work (Riebli et al., 2013) described a set of embryonically born (i.e., primary) neurons belonging to the DM1-4 lineages. These neurons, visualized by the expression of R45F08-Gal4 (Jenett et al., 2012), form a commissural tract that becomes incorporated into the fan-shaped body. Along with the emerging tracts and filopodia extended by secondary neurons of DM1-4 the R45F08-Gal4-positive neurons form a “fan-shaped body primordium” (prFB; Riebli et al., 2013; Hartenstein et al., 2015). In the present paper, we undertook a detailed analysis of the structure, differentiative fate, and developmental role of the primary neurons that form the fan-shaped body primordium, using serial electron microscopy of the early larval brain in combination with confocal analysis of all stages covering early larva to adult. We show that the neurons of the fan-shaped body primordium, that we will call fan-shaped body pioneers (FB pioneers) in the following, form part of a much larger population of early larval brain neurons that are arrested in development, projecting a simple, thin, unbranched process into the neuropil. The large majority of these neurons entirely lack synaptic contacts; in a small number of them, a few presynaptic sites are seen.

Aside from their undifferentiated neurite arbor, these neurons differ from regular, mature neurons by the small size of their cytoplasm and nucleus, and the abundance of heterochromatin. Virtually every brain lineage possesses a complement of the small undifferentiated (SU) neurons. Our data show further that SU neurons differentiate in the late larva and pupa and give rise to distinct adult neuron populations; FB pioneers produce the pontine neurons of the fan-shaped body. Later born secondary neurons of the DM1-4 lineages, destined to form the various classes of unicolumnar neurons of the central complex, fasciculate with the FB pioneers on their pathway towards and across the midline. However, selective ablation of FB pioneers did not result in gross abnormalities of the trajectories of unicolumnar neurons, suggesting that the initial axonal pathfinding of the two system of columnar neurons may be controlled independently.

## Results

### Embryonically born neurons arrested prior to terminal differentiation prefigure adult axon tracts

Based on size and chromatin structure, four types of neuronal cell bodies can be distinguished in the brain. The most frequent type is represented by cell bodies of differentiated primary neurons. These cells measure 12-24*µ*m^*2*^in cross sectional area; their cytoplasm is relatively voluminous and electron light (Fig.2A, C). Each cell body projects a single process towards the neuropil. Processes of neurons of one lineage collect into cohesive bundles, the primary axon tracts (PATs), before entering the neuropil (Fig.2B; see below). Primary axon tracts continue for variable distances in the neuropil, before splitting up into individual, branched axon trees forming synapses. The second type of cell body belongs to mitotically quiescent neuroblasts. Resembling differentiated neurons in size and shape, neuroblasts can be distinguished from these cells by an electron dense cytoplasm and chromatin rich nucleus (Fig.2A). Quiescent neuroblasts are also connected to the neuropil by a single fiber that follows the PAT of its lineage (Lovick et al., 2016); however, unlike differentiated neuronal axons, these neuroblast-derived fibres end near the neuropil surface.

**Figure 2.**
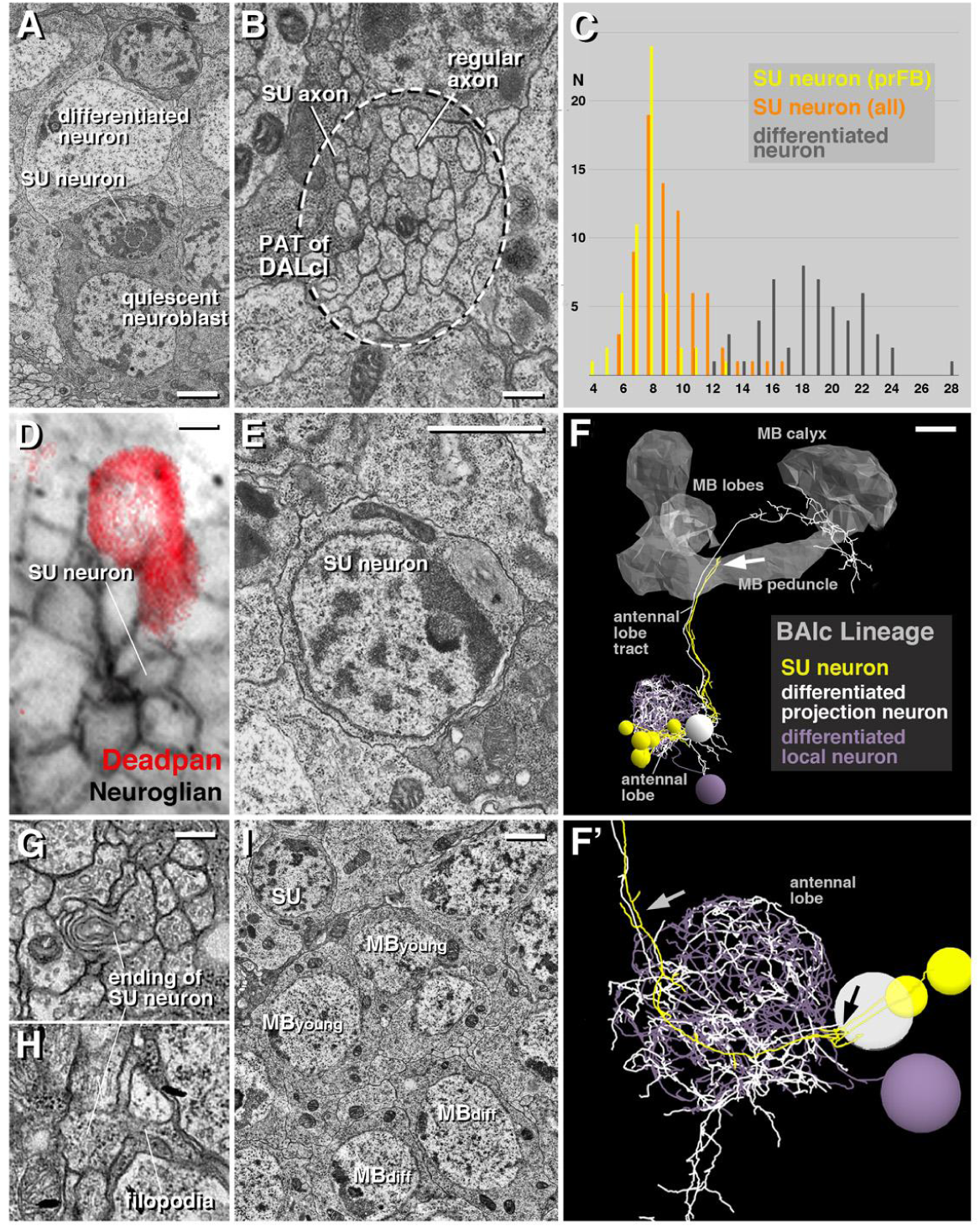
Structural features of small undifferentiated (SU) neurons in the early larval brain. (A) Electron micrograph of brain cortex illustrating neural cell bodies, including differentiated neuron (top), SU neuron (middle) and quiescent neuroblast (bottom). (B) Electron micrograph of cortex-neuropil boundary, showing bundle of axons (surrounded by hatched line) formed by the axons of the two DALcl lineages as they enter the neuropil. Within this bundle, axons of small undifferentiated neurons (“SU axon”) form coherent contingent of thin, electron dense fibers; they are surrounded by thicker and typically more electron-lucent axons of differentiated neurons (“regular axon”). (C) Histogram showing size distribution of differentiated neurons (grey; number counted =) and SU neurons (orange: all SU neurons of right brain hemisphere; n =; yellow: SU neurons forming fan-shaped body primordium (prFB); n =). Units on horizontal axis are in *µ*m2 and represent area of cut surface of cell body. (D) Confocal section of brain cortex of first instar larva. Antibody against Neuroglian labels primary neurons (white); anti-Deadpan labels neuroblasts which at this stage start to enlarge and divide. Note clusters of small cell bodies (SU neuron) interspersed among large, differentiated cells. (E) Electron micrograph of cell body of SU neuron, featuring abundant heterochromatin and extremely thin rim of cytoplasm surrounding round nucleus. (F, F’) 3D digital model of part of one primary lineage (BAlc), illustrating structural aspects of SU neurons. BAlc includes local interneurons and projection neurons connecting the antennal lobe with higher brain centers (Berck et al., 2016). Shown are one differentiated projection neuron (mPN iACT A1 white) and one differentiated local interneuron (Broad D1:grey) whose dendritic arborizations outline the antennal lobe; the axon of the projection neuron follows the antennal lobe tract. Shown in yellow are the SU neurons which were identi1ed for BAlc. They each form an unbranched axon that follows the axons of the differentiated neurons as they enter the antennal lobe neuropil (black arrow in F’) and extends into the antennal lobe tract (grey arrow in F’) where they end approximately midway (white arrow in F). The model is presented in lateral view; anterior to the left, dorsal up. The mushroom body (MB) is outlined in grey for spatial reference. (G, H) Electron micrographs show club-shaped endings of SU axons, featuring membrane lamellae (G) and short filopodia (H). (I) Electron micrograph of mushroom body cortex. Shown are two cell bodies of differentiated neurons (MBdiff) located next to two young neurons (MByoung). These show similar size and chromatin structure as differentiated neurons. Bars: 1 *µ*m (A, E, I), 0.25 *µ*m (B, G, H), 2 *µ*m (D), 5 *µ*m (F).

A third type of neural cell body is characterized by its small size, scanty, electron dense cytoplasm, and heterochromatin-rich nucleus (Fig.2A, E). These small neurons, measuring 4-16*µ*m^*2*^in cross sectional area, project 1bers which are signi1cantly thinner and more electron-dense than the axons of differentiated neurons (Fig.2B). They follow differentiated axons into the neuropil, staying together as a tight bundle, without branching or forming synaptic contacts (Fig.2F, F’). Dense axons end abruptly with club-shaped endings, often producing short 1lopodia (Fig.2G, H). We interpret the small, heterochromatin-rich neurons, which (based on size) are also visible light microscopically (Fig.2D) as late born neurons which fail to differentiate in the embryo, and therefore call them “small undifferentiated (SU)” neurons in the following.

The fourth type of neural cell is the actively dividing neuroblast, of which the 1rst instar brain contains 1ve. Four of these neuroblasts are associated with the mushroom body (MB); a 1fth one belongs to lineage BAlc. Active neuroblasts are much larger than quiescent ones; they are located at the brain surface, and do not possess a process towards the neuropil (not shown). Continued proliferation of the active MB neuroblasts results in a steady increase in the number of MB neurons; thus, unlike other lineages, production of MB neurons is not interrupted by a quiescent period to form a discrete embryonic (primary) phase, separated from a larval (secondary) phase. As a result, the MB neurons reconstructed in a recent publication (Eichler et al., 2017) show different degrees of structural maturity, from fully differentiated neurons over “young” neurons with fewer branches and synaptic contacts to “very young” neurons that resemble SU neurons, having a short, unbranched 1ber that barely reaches the peduncle. Interestingly, the cell body of young and very young MB neurons does not differ in size or chromatin structure from that of fully differentiated neurons (Fig.2I). This 1nding supports the idea that young MB neurons, unlike the SU neurons of other lineages, do not enter a prolonged phase where differentiation is arrested; instead, they go directly into differentiation mode and become integrated into a growing mushroom body circuitry. SU neurons are found scattered throughout the brain, but with higher density in the dorsomedial area, where they form large, coherent clusters (Fig.3A-C).

**Figure 3.**
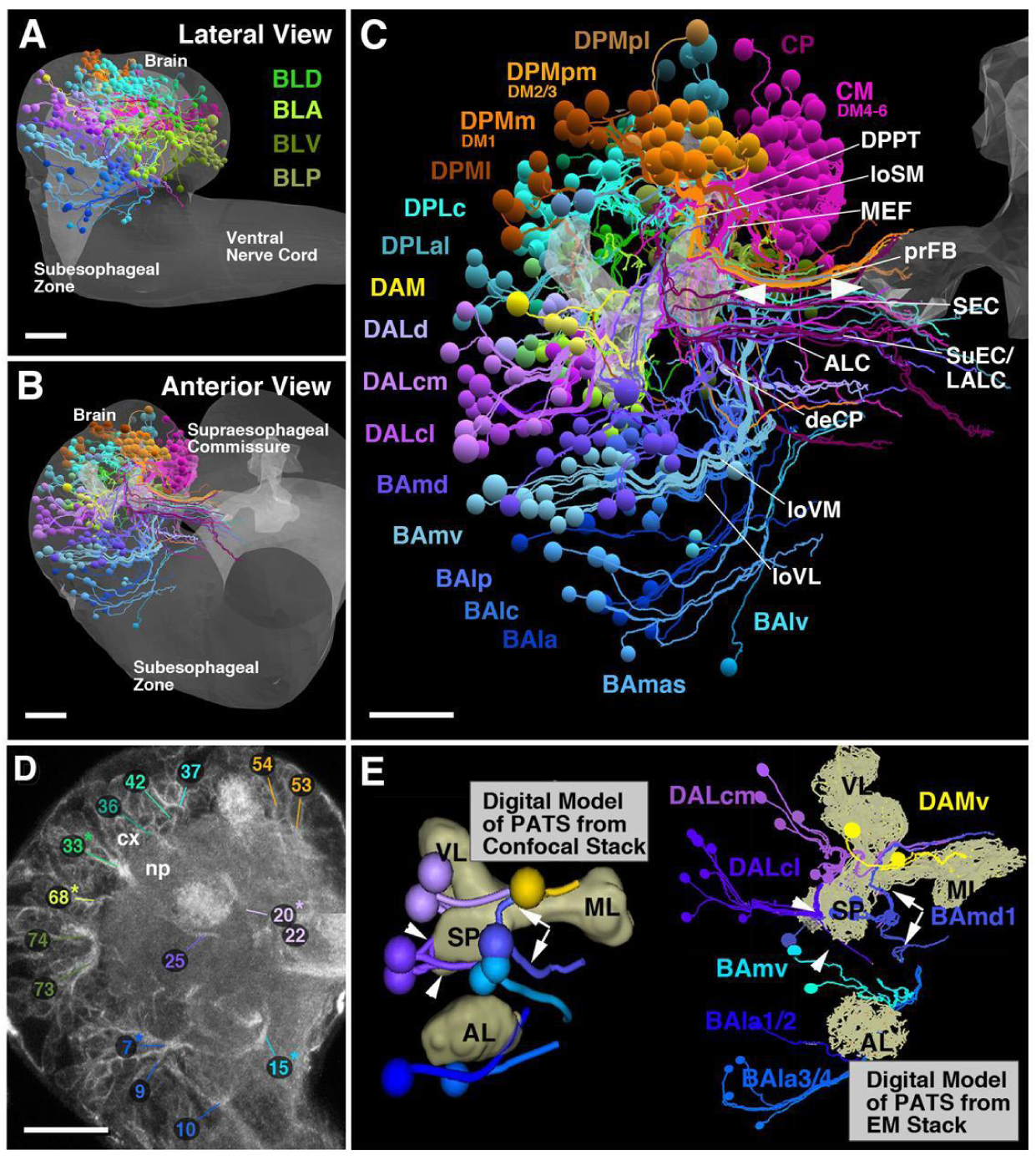
Pattern of SU neurons and primary axon tracts (PATs) in the early larval brain. (A-C) 3D digital model of first instar larval brain in lateral view (A; anterior to the left, dorsal up) and anterior view (B, C; dorsal up). Cell bodies and axons of all SU neurons of one hemisphere are shown as colored spheres and lines. Differential coloring indicates association of SU neurons with spatially contiguous groups of lineages; labels of lineages are shown in corresponding colors (A, B; for nomenclature of lineages see Hartenstein et al., 2015). In (A, B), outline of brain and ventral nerve cord is rendered in grey; in (C), outlines of mushroom body lobes, located in center of neuropil, are shaded, with tips of left and right medial lobe pointed out by arrowheads. Note that lineage-associated bundles of SU axons follow discrete fascicles identified for the brain throughout development. Among the examples made explicit are the BAmv axons that form the longitudinal ventromedial fascicle (loVM); axons of BAlp constitute the longitudinal ventro-lateral fascicle (loVL); DALd axons form the central protocerebral descending fascicle (deCP); CM lineages the medial equatorial fascicle (MEF); DPMpm the longitudinal superior-medial fascicle (loSM) and primordium of the fan-shaped body (prFB). (D, E) Primary lineage tracts imaged light microscopically can be identified with electron microscopically reconstructed SU axon tracts. (D) shows representative frontal confocal section of early larval brain; primary neurons and their axon tracts are labeled by anti-Neuroglian (white). Primary axons converge at the cortex (cx)-neuropil (np) boundary at invariant positions to form tracts with characteristic trajectories. A subset of tracts, reconstructed from a confocal stack of an anti-Neuroglian labeled brain hemisphere, is shown on the left side of panel (E). The mushroom body lobes (ML medial lobe; SP spur; VL vertical lobe) and antennal lobe (AL) is shown for reference. Right side of (E) shows groups of SU neurons whose axon bundles can be identified with the lineage tracts on left side; note for example axon tract of DALcl which approaches the spur from laterally, and then bifurcates to continue dorsally and ventrally of the spur (white arrowheads). The example pointed out by white arrows represents the tract of BAmd1 which approaches the medial lobe from anteriorly and then bifurcates into a dorsal and ventral branch; both branches turn medially to form distinct commissural tracts. For all abbreviations see Table 1. Bars: 20*µ*m type II lineage, CP2 (DL1); the other two originate as part of the lineage DPMm2 (Fig.4E, F).

**Figure 4.**
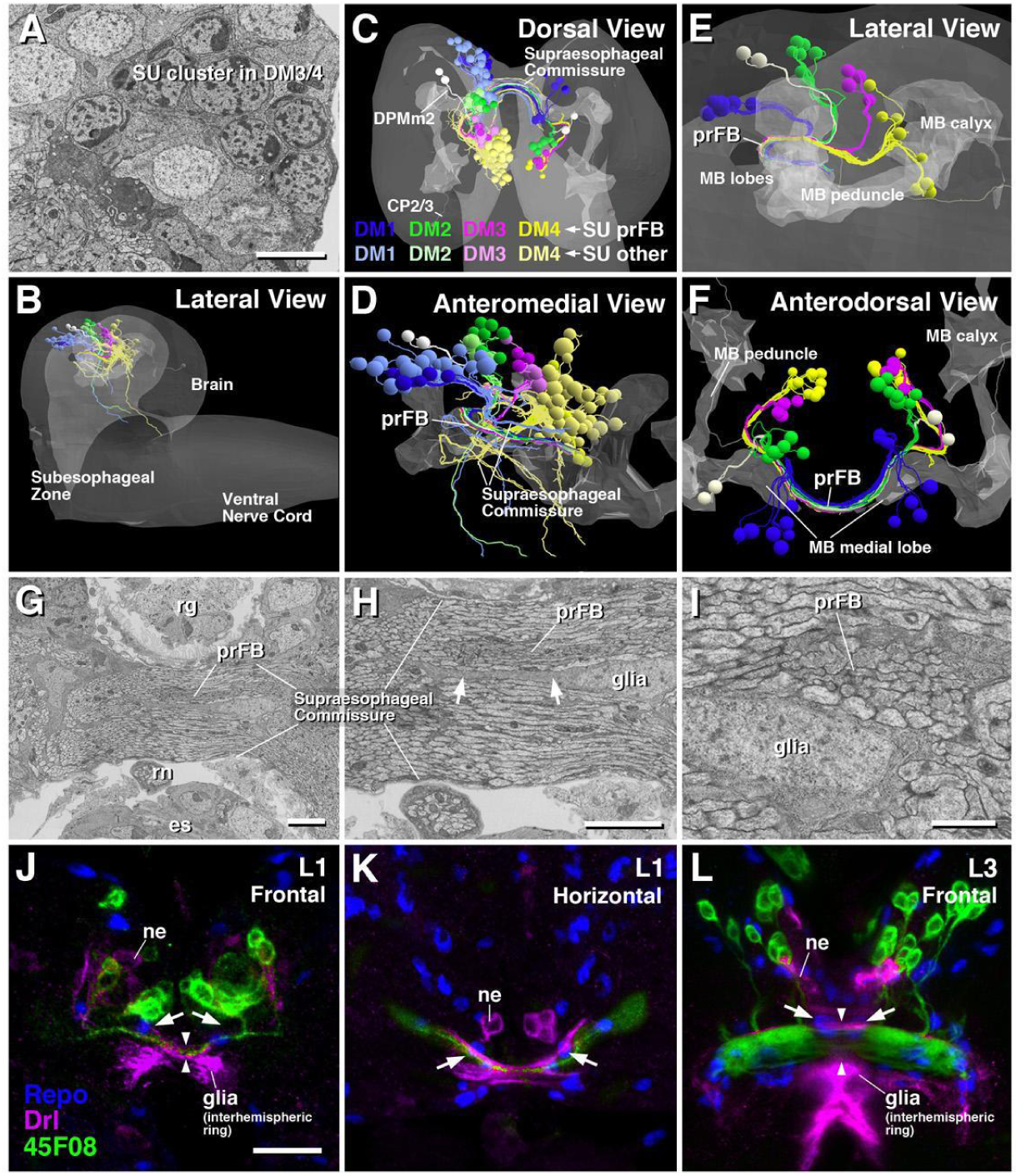
SU neurons of lineages DM1-4 form the primordium of the fan-shaped body (prFB). (A) Electron micrograph of dorso-medial brain cortex showing clusters of contiguous SU neurons associated with lineages DM1-4. (B-F) Digital 3D models of first instar brains showing SU neurons associated with DM1-4 rendered in different colors (DM1 blue; DM2 green; DM3 magenta; DM4 yellow). In all models, outline of mushroom body (representing center of brain neuropil) is rendered in light gray; in (B, C, E), outline of brain is shown in dark gray. In (B-D), all SU neurons of DM1-4 are included in models; subsets of SU neurons contributing to the prFB are shown in saturated colors, other SU neurons are rendered in light colors. In (E, F), only prFB-associated SU neurons are shown. (G-I) Electron micrographs of brain midline with supraesophageal commissure. The prFB appears as a bundle of electron-dense axons (G, H) embedded in fascicles of lighter axons belonging to differentiated neurons. Glial lamella flanks the prFB (arrows in H). Note conspicuous, medially located nucleus of glial cell right adjacent to the prFB. (J-L) Z-projections of frontal (J, L) or horizontal (K) confocal sections of first instar brain (J, K) and late third instar brain (L). Glial nuclei are labeled by anti-Repo (blue); glia surrounding supraesophageal commissure (=interhemispheric ring glia) is labeled by Drl-lacZ (magenta); neurons of prFB are marked by driver line R45F08-Gal4>UAS-mcd8GFP (green). Interhemispheric ring glia forms several channels containing individual commissural fascicles. One channel contains the R45F08-Gal4-positive axon bundle constituting the prFB [arrowheads in (J, L)]. This channel widens as prFB gains in volume towards late larval stages (L). Note pair of large glial nuclei (arrows) located near the midline, posteriorly adjacent to the prFB. These cells correspond in shape, size and position to the prFB-associated glial cells that appear in electron micrographs (see panels H, I). Other abbreviations: es esophagus; ne Drl-lacZ-positive neurons; rn recurrent nerve; rg ring gland. Bars: 2*µ*m (A, G, H); 1*µ*m (I); 10*µ*m (J-L)

**Table 1.**
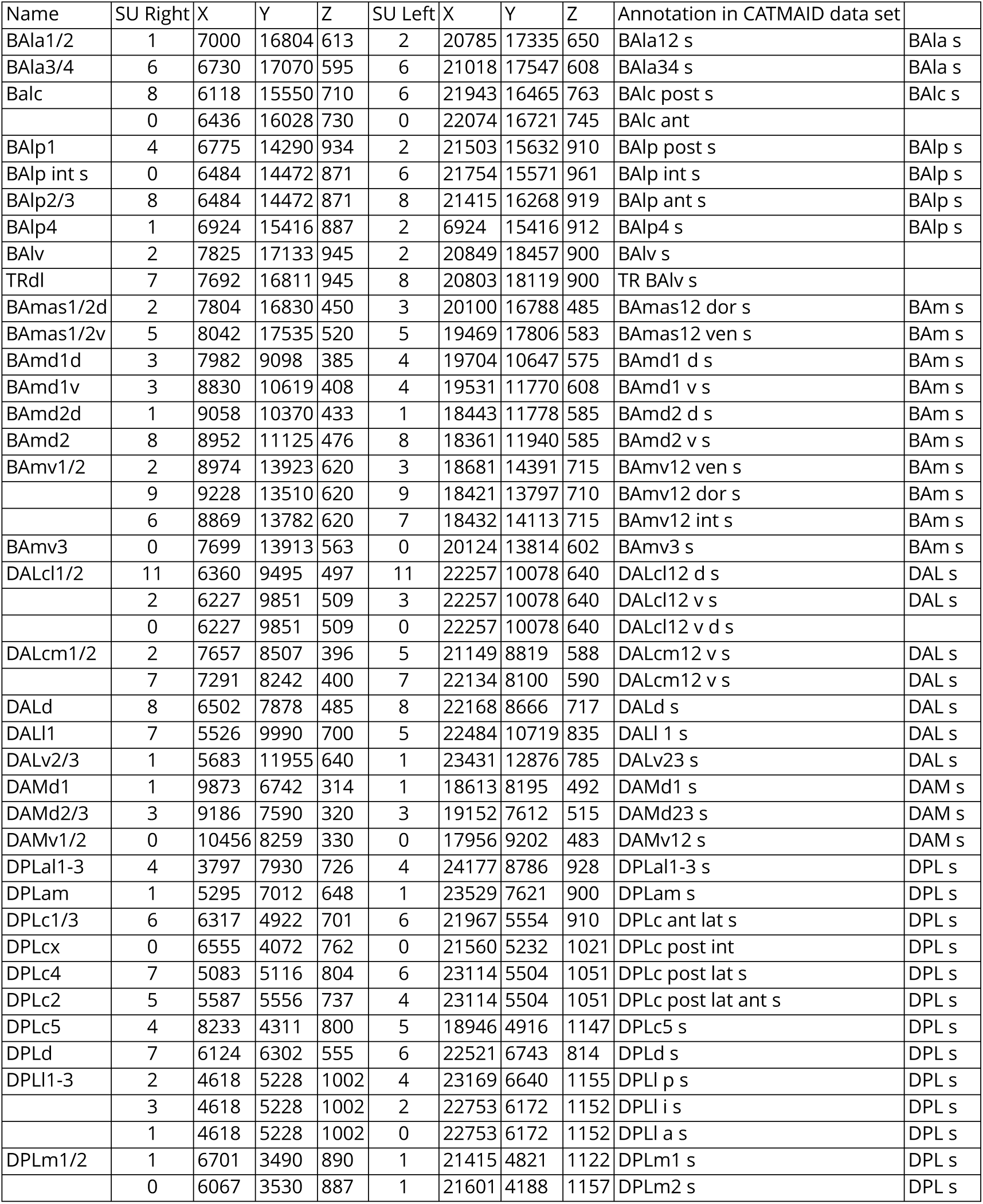
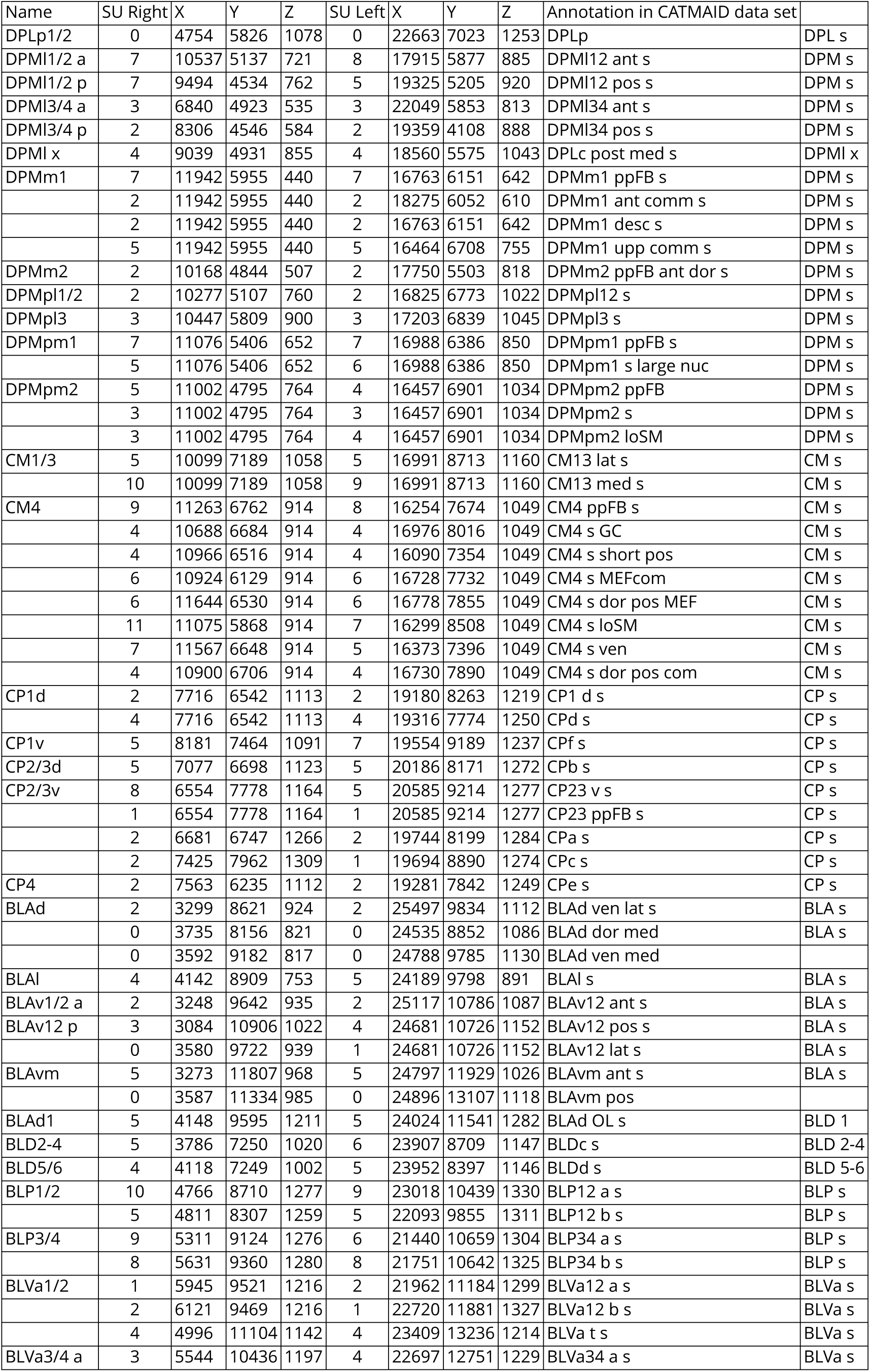
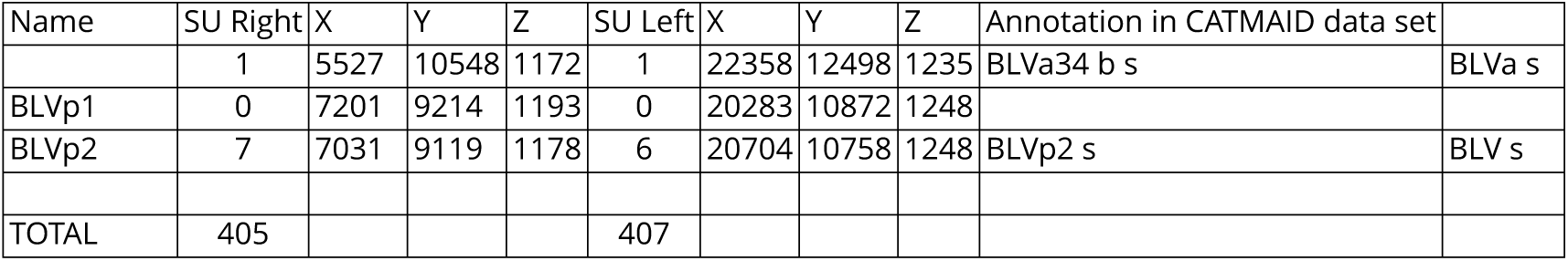
Tabular representation of all SU neurons of the 1rst instar (L1) larval brain ordered by lineages (see also supplementary Figures S2–S9 for graphic representation). SU neurons of a given lineage are clustered in a cohesive tract, the primary axon tract. These tracts provide useful landmarks for the ongoing analyses of the L1 brain circuitry using the CATMAID L1 serial TEM dataset. Column to the left lists names of lineages or lineage pairs/small lineage groups (see Hartenstein et al., 2015). Columns 2 and 6 give numbers of SU neurons associated with corresponding lineages of the right hemisphere and left hemisphere, respectively. Columns 3-5 and 7-9 show the coordinates of the neuropil entry points of the SU-de1ned primary axon tracts for the right hemisphere and left hemisphere, respectively. The origin (z=0/x=0/y=0) represents the anterior/right/dorsal corner of the virtual cuboid that encloses the data set. The units for z is section number (z=0 corresponding to most anterior section; section thickness = 45nm); x and y indicate pixels, with each pixel corresponding to approximately 3.8nm. The two columns to the right show how SU neuron populations are currently annotated in the CATMAID data stack..

SU neurons amount to a signi1cant fraction (more than 25%) of the overall number of primary neurons of the brain (Table 1). Almost every lineage-associated primary axon tract possesses between one and more than 10 of SU axons (Fig.3C; Table 1). Comparison between left and right brain hemisphere showed that the exact number of SU neurons accompanying lineages is relatively invariant (Table 1).

Axons of SU neurons of a given lineage typically form a tight sub-bundle, often near the center of a given PAT (Fig.2B). The trajectories of these SU tracts in all cases matched the pathway followed by primary neurons, as exempli1ed for the lineage BAlc, forming the antennal lobe tract, in Fig.2F. Based on these trajectories we were able to identify the PATs reconstructed from the L1 EM stack with lineages de1ned for the 1rst instar larval brain in previous works (Hartenstein et al., 2015). The 1rst instar larval brain features more than 50 discrete PAT 1ber bundles with characteristic neuropil entry points and trajectories (Fig.3D). Once digitally reconstructed with CATMAID software; (Saalfeld et al. 2009, Schneider-Mizell et al. 2016) and displayed in the 3D viewer, PAT 1ber bundles can be compared with those in the light microscopic 1rst instar brain map (Fig.3E), and thus identi1ed as belonging to speci1c lineages. Note, as examples shown in Fig.3E, characteristic pattern elements visible in confocal image and EM stack, such as the branching of the DALcl1/2 tract around lateral surface of spur (a domain within the mushroom body), or the BAmd1 tract branching anterior to the mushroom body medial lobe. The neuropil entry points and trajectories of SU neuronal tracts provide useful landmarks for the ongoing analyses of the 1rst instar larval brain circuitry using the 0111-8 L1 ssTEM volume (Ohyama et al. 2015); both sets of data are presented graphically and numerically in supplementary figures S2-S9.

### SU neurons of the four lineages, DM1-DM4, form a glia ensheathed fan-shaped body primordium in the 1rst instar larva

In the dorso-medial domain of the brain, which houses all lineages later giving rise to the columnar neurons of the central complex (the type II lineages DM1-DM4; Bello et al., 2008), SU neurons form large clusters of cells (Fig.4A). Based on axonal trajectory we were able to recognize the individual DM lineages and their associated SU neurons. DM1-4 are aligned along the brain midline, with DM1 occupying the most anterior, and DM4 the most posterior position (Fig.4B, C). For each of the DM lineages, SU neurons form several subpopulations with axons following different pathways (Fig.4D). Axons of one SU subpopulation per DM lineage, containing between 5 and 7 neurons, converge on a shared commissural bundle that crosses the brain midline in the center of the supraesophageal commissure (Fig.4E, F). This bundle stands out from surrounding tracts by its higher electron density, being composed almost entirely of thin SU axons (Fig.4G, H); furthermore, two paired glial cells ensheath the bundle on all sides (Fig.4H, I). Aside from the subpopulations of SU neurons of DM1-4, three other SU axons join the bundle. One of these is derived from another type II lineage, CP2 (DL1); the other two originate as part of the lineage DPMm2 (Fig.4E, F).

For late larvae, the *pointed* enhancer fragment Gal4-driver R45F08-Gal4 has been reported to be expressed in small subpopulations of neurons of DM lineages whose axons form a glia-covered commissural structure that will enlarge into the fan-shaped body, and was therefore called fan-shaped body primordium (prFB; Riebli et al., 2013). In 1rst instar larvae, R45F08-Gal4 highlights four paired clusters of neurons in the dorso-medial brain whose axons form a thin tract that, in regard to size, central location within the brain commissure, and glial enclosure corresponds to the structure identi1ed electron microscopically (Fig.4J-L). We conclude that the fan-shaped body primordium is formed in the embryo by four paired sets of small undifferentiated neurons which we will call “fan-shaped body pioneers” (FB pioneers) in the following.

### FB pioneers are topologically ordered and give rise to the pontine neurons of the fan-shaped body

FB pioneer neurons exhibit a strict topological order with regard to the position and average length of axons. Axons of neurons of a given DM lineage remain together as a bundle throughout the length of the prFB (Fig.5A-C). The DM1 bundle enters anteriorly and dorsally, and remains at the dorsal edge of the prFB, followed by DM2, DM3 and DM4, which is located most ventrally (Fig.5C-E). DM4 axons are the shortest, most of them ending before reaching the midline (Fig.5B, J). Axons of DM1-DM3 cross the midline but extend across for various distances, in correlation to their lineage identity: DM1 axons cross furthest, followed by DM2 and DM3 (Fig.5B, J).

**Figure 5.**
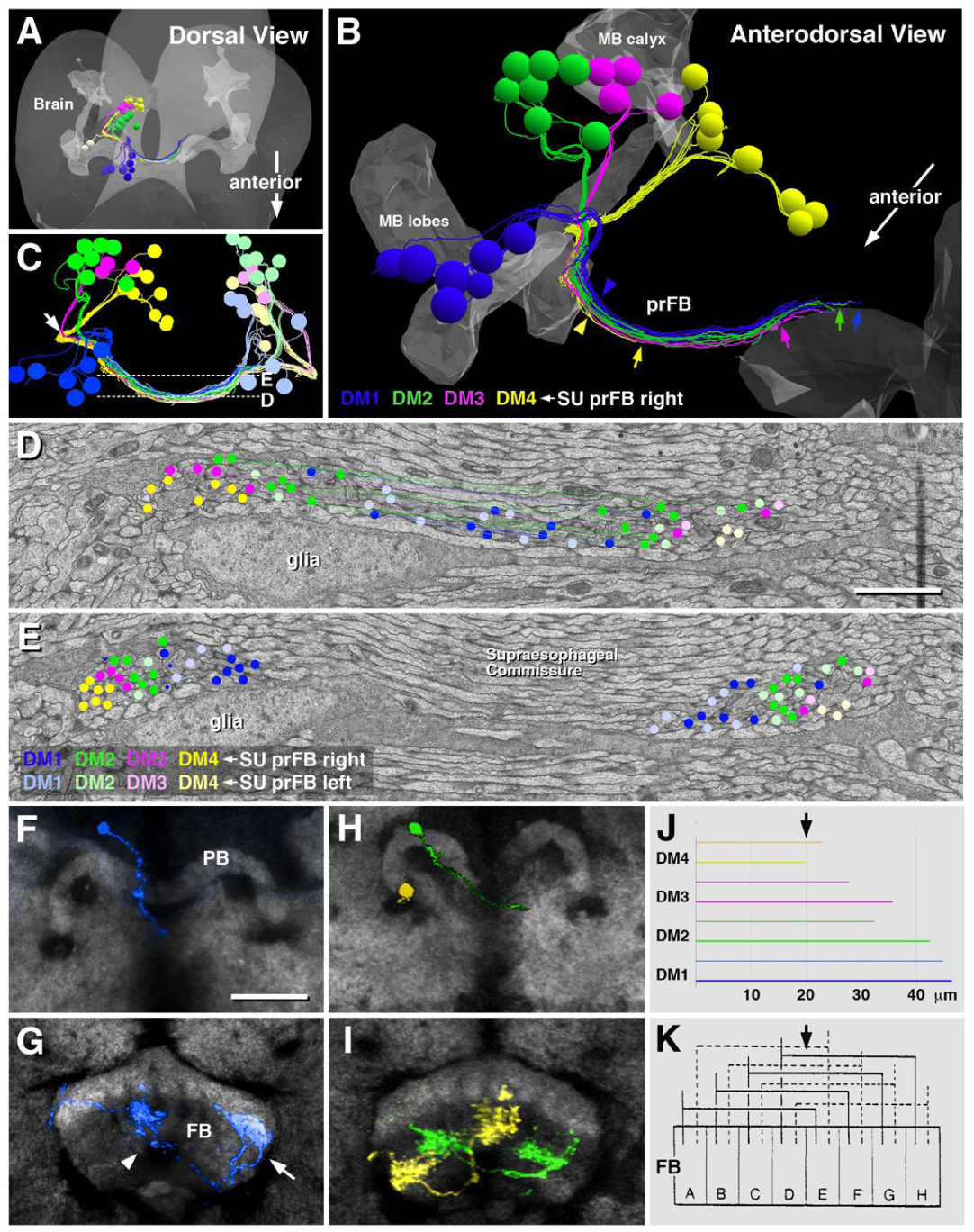
Topological order and developmental fate of FB pioneers. (A-C) Digital 3D model of the EM larval brain showing FB pioneers rendered in different colors. The mushroom body (MB) is rendered in gray for reference. (A) and (B) show FB pioneers of the right hemisphere; in (C), right hemispheric FB pioneers appear in saturated colors, left hemispheric ones in light colors. Note relationship between lineage identity and axonal projection among FB pioneers. Cells of DM1 (blue), located furthest anterior in the brain, have axons located at the dorso-posterior edge of the prFB (blue arrowhead in panel B), and extend furthest in the contralateral hemisphere (blue arrow in B). DM4 axons (yellow) originate furthest posteriorly in the brain (A, B), are located most ventro-anteriorly in the prFB (yellow arrowhead in B) and are the shortest, barely crossing the midline (yellow arrow in B). FB pioneers of DM2 (green) and DM3 (magenta) fall in between these extremes. (D, E) Electron micrographs of prFB sectioned at the two planes shown by hatched lines in panel (C). Sectioned profiles of FB pioneers were assigned to their cell bodies of origin and are demarcated by circles, color coded following the same scheme used in the 3D digital models in (A-C). Axons belonging to a lineage form a coherent bundle within the prFB. Note that bundles of corresponding lineages of the left and right hemisphere project together (light blue circles are closest to saturated blue circles, etc). (F-I) Frontal confocal sections of central complex of adult brain, showing single cell clones of neurons descended from FB pioneers. (F, H) are sections at the level of the protocerebral bridge (PB), (G, I) at the level of the fan-shaped body (FB). Labeled clones (for technique, see Material and Methods) are color coded following the same scheme used in panels (A-E). (F, G) show a pontine neuron of lineage DM1 (blue), with a medially located cell body and axon entering at the medial PB (F), and terminal axons branching in an ipsilateral medial column (arrowhead in G) and a contralateral lateral column (arrow in G). (H, I) show a DM4 clone (yellow) and DM2 clone (green). (J) Histogram depicting average length of FB pioneer axons belonging to lineages DM1-4. Upper horizontal bar of each pair belongs to left hemispheric lineage, lower bar to corresponding right hemispheric lineage. The sharp medial turn of axon as it enters the prFB (white arrow in panel C) was taken as origin on x-axis; arrow at top demarcates brain midline. (K) Schematic of Hanesch et al. (1989), showing fan-shaped body (FB) divided into eight columns (A-H; arrow at top indicates midline). Based on Golgi-preparations, Hanesch et al. distinguished between four classes of bi-columnar pontine neurons, indicated by solid lines on left side, and hatched lines on right side. Our data indicate that the four classes described by Hanesch et al. correspond to the four lineages of origin of the pontine neurons. Bars: 24*µ*m (D, E); 25*µ*m (F-I)

Using the R45F08-Gal4 driver we followed the prFB neurons through larval and pupal development into the adult (see below), and generated single cell MCFO clones (Nern et al., 2015) to visualize their morphology as differentiated cells. All neurons forming the early larval prFB give rise to pontine neurons, recognizable by their regular bi-columnar organization within the fan-shaped body (Fig.5F-I). Labeled neurons match the four-fold symmetric scheme elaborated by Hanesch et al. (1989) for pontine neurons, whereby neurons entering the FB most laterally (DM1; Yellow in Fig.5H, I) branch near their point of entry in the lateral column “A”, then cross the midline, and have their second arborization in a column right next to the midline (“E”; Fig.5K, from Hanesch et al., 1989). Neurons of DM1 (blue in Fig.5F, G), as well as DM3 and DM4 (green and yellow, respectively, in Fig.5H, I) have a similar structure, innervating staggered pairs of columns, with one column always ipsilaterally relative to the location of cell bodies, and one contralaterally (Fig.5K). Note that the length distribution of SU axons of DM1-DM4 coincides with the adult pattern already in the early larva: DM4 axons are shortest, barely reaching the midline; DM1 axons are longest (compare Fig.5J, K)

### Later born columnar neurons of DM1-DM4 follow the trajectory prefigured by the FB pioneers

R45F08-Gal4-positive axons of FB pioneers can be followed throughout larval and pupal development into the adult stage. Until 48h after hatching (L48), the prFB formed by these axons remains as a single axon bundle located in the center of the brain commissure (Fig.6A). At this stage, the secondary phase of neuron production has commenced (Lovick and Hartenstein, 2015). DM1-DM4 form secondary axon tracts, labeled by anti-Neurotactin, that grow along the R45F08-Gal4-positive (and neurotactin-negative) FB pioneers (Fig.6B, B’, C, C’).

**Figure 6.**
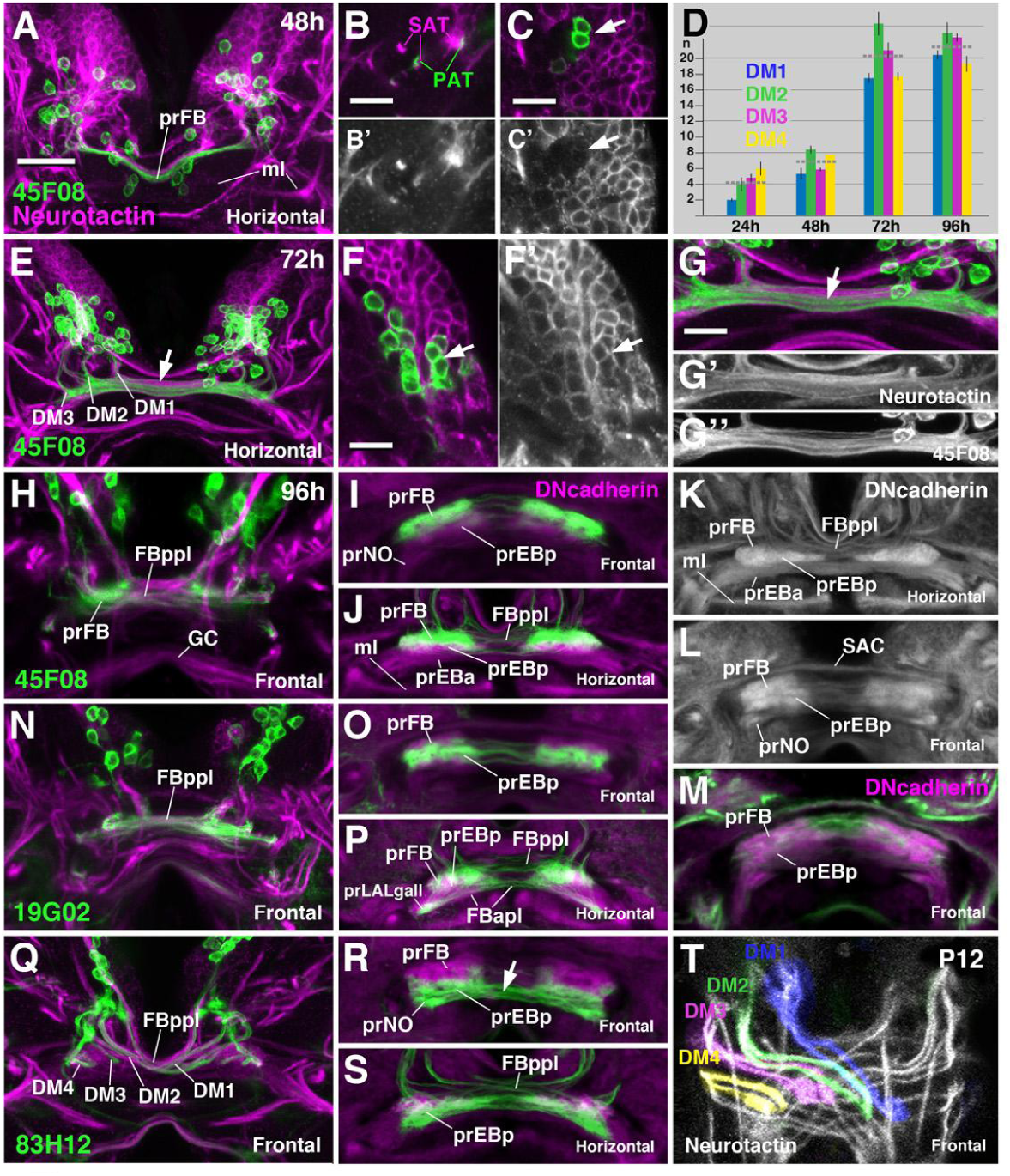
Growth and development of the prFB during the larval period. Panels show Z projections of frontal or horizontal confocal sections of larval brains at different stages (48h, 72h, 96h after hatching). Global labeling of secondary (larvally born) neurons with anti-Neurotactin (magenta in A-C, E-G, H, N, Q; white in B’, C’, F’, G’, T) or neuropil with anti-DNcadherin (magenta in I, J, M, O, P, R, S; white in K, L). FB pioneers are labeled by R45F08-Gal4>mcd8UAS-GFP (green in A-C, E-J). (A-C) At 48h (second larval instar) FB pioneers form a single bundle as in freshly hatched first instar (see Fig.4J, K). The outgrowing secondary axons tracts (SATs) of DM1-4 (magenta in B) fasciculate with FB pioneers (green in B). At this stage, R45F08-Gal4 expression is still confined to Neurotactin-negative primary FB pioneers (C, C’). (D) Average numbers and standard deviations of R45F08-Gal4-positive DM1-4 neurons between 24 and 96h after hatching (n=5). Horizontal hatched lines give average for all four DM lineages. (E-G”) At 72h after hatching (early third instar) the FB pioneer axons have split into several commissural bundles (arrow in E, G). Gaps between these bundles are filled with Neurotactin-positive secondary fibers formed by lineages DM1-4 (G-G”). Numerous Neurotactin-positive secondary neurons have joined the set of R45F08-Gal4-positive FB pioneers (arrow in F, F’). (H-L) At 96h after hatching (late third instar) secondary DM1-4 axons have increased in number and form a system of crossing bundles, the posterior plexus of the fan-shaped body (FBppl in H, J, K). Both R45F08-Gal4-positive FB pioneers and secondary axons display tufts of filopodia which appear as DNcadherin-positive domains (prFB, prEBp, prEBa in panels I-L). (M-P) Labeling of secondary PB-FB-LALgall and PB-EB-LALgall neurons by R19G02-Gal4 driver. Terminal filopodia of these neurons are found in ventral prFB, prEBp and prLALgall (see also Lovick et al., 2017). (Q-S) R83H12-Gal4 is expressed in a different subset of secondary neurons, mainly PB-FB-NO and PB-EB-NO. Note in (Q) chiasmatic architecture of projection of labeled neurons towards the fan-shaped body, with all four bundles turning medially, but only DM1 and DM2 crossing towards contralaterally, and DM3/4 remaining ipsilaterally. This chiasmatic projection is typical for all unicolumnar neurons, as demonstrated in (T; 12h after puparium formation), where anti-Neurotactin (white) globally labels massive bundles of secondary axons of DM1-4. Bar: 25*µ*m (for all panels except B-C’, F-G”); 10*µ*m (for B-C’, F-G”).

The size and appearance of the prFB changes at later larval stages. Around 72h after hatching, the number of neurons labeled by R45F08-Gal4 has increased over that counted at 24h (Fig.6D, E), and many of these newly R45F08-expressing neurons form part of the Neurotactin-positive clusters of secondary neurons (Fig.6F-G’). In addition, a notable change in the structure of the prFB has taken place, whereby the initially coherent tract of primary FB pioneers has split into several thinner bundles (Fig.6G, G”). The spaces in between these is 1lled by the secondary, Neurotactin-positive axons of DM1-4 (Fig.6G’), which give rise to the uni-columnar neurons of the central complex. Several markers for the major classes of uni-columnar neurons exist, among them the Gal4-drivers R9D11-Gal4 Jenett et al., 2012; Riebli et al., 2013), R19B02-Gal4 (Jenett et al., 2012; Wolf et al., 2015; Omoto et al., 2017), and R83B12-Gal4 (Jenett et al., 2012; Wolff et al., 2015; Omoto et al., 2017; this study). R19B02 is expressed in several classes of neurons connecting PB, dorsal FB, and EB with the neuropil laterally adjacent to the central complex (PB-EB-gall and PB-FB-rub neurons of Wolff et al., 2015; supplementary Fig.S1B); R83H12 is expressed in neurons connecting the PB, ventral FB, EB and the NO (PB-FB-NO and PB-EB-NO neurons of Wolff et al., 2015; supplementary Fig.S1C). In conjunction with R45F08-Gal4, these markers allowed us to discern the distribution of pontine versus uni-columnar neurons in the prFB of the late larva, as presented below.

In the late larva, the fan-shaped body primordium occupies a sizeable volume as a result of the ingrowth of secondary axons, in conjunction with the formation of tufts of 1lopodia that pre1gure the fan-shaped body and noduli. Primary and secondary axon tracts of DM1-DM4 form a plexus of fiber bundles at the posterior surface of the fan-shaped body primordium (posterior plexus of the fan-shaped body (FBppl) in Fig.6H, J, K). Within this plexus, the R45F08-Gal4-positive pontine neuron precursors appear as thin axon bundles crossing the midline (Fig.6H, J). Also secondary tracts of uni-glomerular neurons, including the 83H12-positive PB-FB/EB-NO neurons and 19G02-positive PB-EB/FB gall neurons, cross in the posterior plexus, forming a system of chiasms that establish the characteristic, modular connectivity of the fan-shaped body and ellipsoid body (Fig.6N, P, Q, S, T). Tracts of DM1 and DM2 cross the midline to terminate in the lateral and medial half of the contralateral prFB, respectively; those of DM3 and DM4 remain ipsilateral, targeting the medial and lateral half of the ipsilateral prFB, respectively (Fig.6Q, T; see also Boyan et al., 2017).

In addition to the posterior plexus of axons, the late larval fan-shaped body contains voluminous tufts of filopodia, visible by their high expression levels of DN-cadherin (Riebli et al., 2013; Omoto et al., 2015; Lovick et al., 2017). These tufts outline three bar-shaped neuropil domains (Fig.6K, L, M). The domain located dorsally and posteriorly will give rise to the neuropil of the fan-shaped body proper (prFB). Accordingly, filopodia of R45F08-Gal4-positive precursors of pontine neurons are restricted to this dorsal compartment (Fig.6I, J). Located further anteriorly and ventrally is the primordium of the posterior ellipsoid body (prEBp; Lovick et al., 2017); the primordia of the noduli (prNO) form two ventrally-directed appendages of the prFB (Fig.6L). 19B02-positive secondary neurons, which in the adult brain branch within the FB as well as the posterior EB, 1ll a volume straddling the boundary between both domains (Fig.6O, P). 83H12-positive filopodia of presumptive PB-EB/FB-NO neurons are concentrated in the most ventral domain of the prFB, the prEBp and the prNO (Fig.6R, S). Axons of these neurons form thick chiasmatic fiber systems that extend in between the ventral surface of the prFB and the prNO of either side (Fig.6R).

### Ablation of the FB pioneers

Given the early appearance of precursors of the pontine system, and its intimate spatial relationship with the subsequently generated unicolumnar neurons, we sought to establish the effect of ablating the former by using the R45F08-Gal4 construct to express UAS-hid;rpr in the FB pioneers. Activating the pro-apoptotic construct between 24h and 48h after hatching by inactivation of temperature-sensitive Gal80ts, eZciently ablated the FB pioneers; it also resulted in a high rate of lethality during larval stages, which is likely due to expression of R45F08-Gal4 in other tissues (V.H., unpublished observation). No late pupa or adult flies under these circumstances survived, whereas non-temperature shifted controls were completely viable. The late larval prFB of the surviving FB pioneer-ablated larvae showed a characteristic structural phenotype. All elements of the prFB described above could still be recognized; however, the DN-cadherin-rich primordia of fan-shaped body and posterior ellipsoid body/noduli were separated by a wider gap (arrowheads in Fig.7B, E, E’). In addition, the posterior plexus of the fan-shaped body, normally formed by a loose assembly of “wavy”, thin bundles (Fig.7A, C), appeared as thick, smooth cable of axons (Fig.7D, F). We conclude that FB pioneers, possibly by establishing an early connection between the left and right half of the prFB through their crossing axons, play a structural role in pulling these halves together into a united structure (Fig.7I, J). Aside from this function, there does not appear to be an effect on the formation and pathway choices of the later born secondary axons: these split into several sub bundles and projected into the prFB in a pattern that showed no gross abnormalities compared to the control (Fig.7G, H). This result suggests that axonal pathfinding of the two systems of small field neurons of the central complex, i.e., the uni- and bi-columnar neurons, is controlled independently.

**Figure 7.**
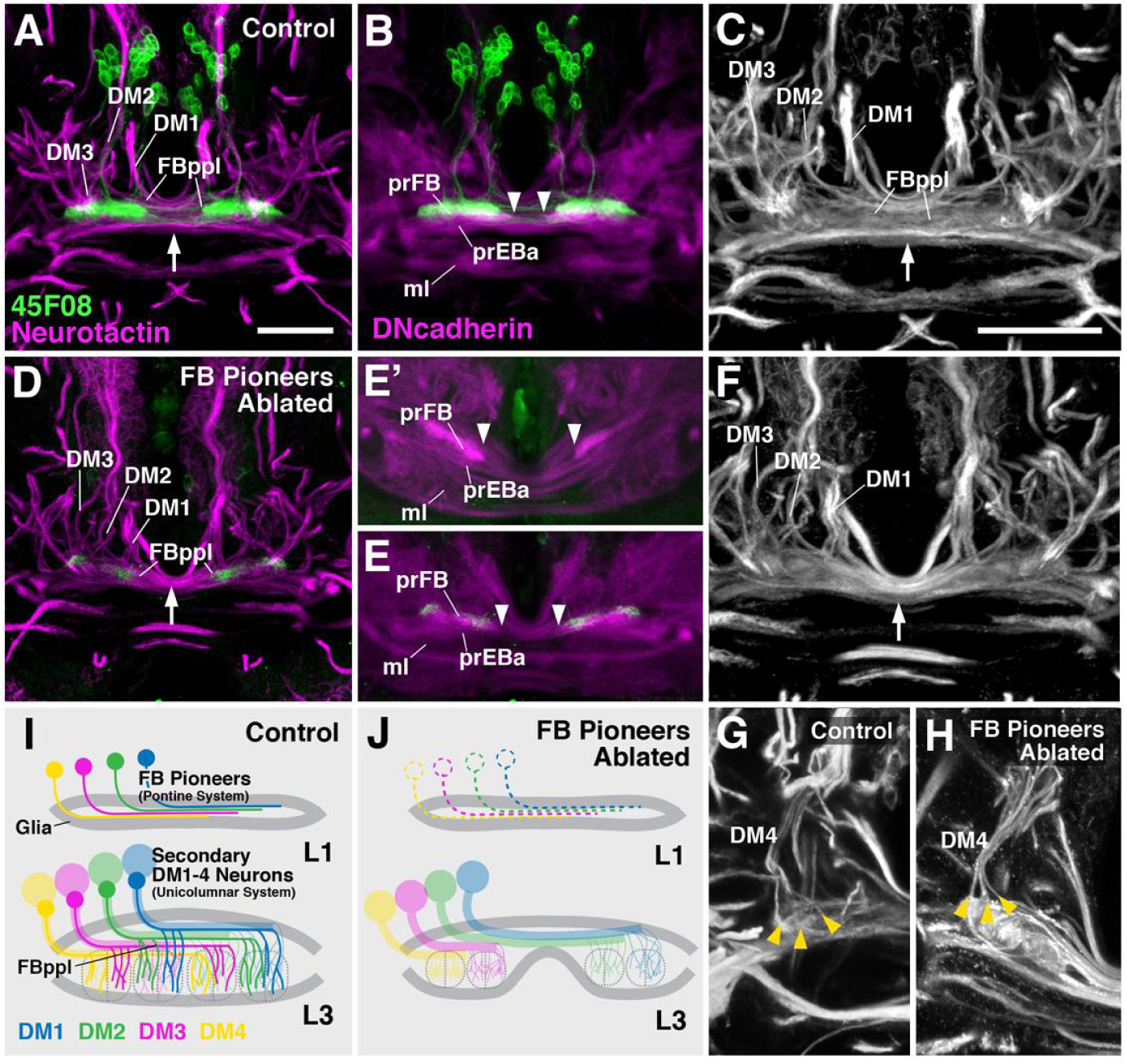
Structure of the prFB after ablation of FB pioneers. (A-C) Z-projections of horizontal confocal sections of wild-type late third instar brains, illustrating entry of lineages DM1-3, posterior plexus of fan-shaped body (FBppl), and filopodial tufts forming primordia of fan-shaped body neuropil (prFB) and ellipsoid body neuropil (prEBa). For a clear depiction of the FB pioneers, males lacking UAS-hid;rpr and tub-Gal80ts are shown. (D-F) Same views of late larval brain as in (A-C) after ablation of FB pioneers by activating UAS-hid;rpr from 24-48h with R45F08-Gal4. (E and E’) show two different representative specimens. Note that FBppl is reduced in diameter (arrow in D, F) compared to control (arrow in A, C). Also primordium of fan-shaped body neuropil (prFB in B, E, E’) is separated by wider gap in ablated specimens (arrowheads in B, E, E’). (G, H) Z-projection of horizontal confocal sections of one brain hemisphere of control (G) and ablated (H) specimen, showing normal branching pattern of secondary axon bundles of DM4 lineage (yellow arrowheads). The neurotactin and DNcadherin pattern of non-temperature shifted female controls (which contain UAS-hid;rpr and tub-Gal80ts), in which no ablation occured, were indistinguishable from A-C, G (not shown). (I, J) Schematic fan-shaped body primordium at first larval instar (L1) and late third instar (L3) in control (I) and ablated specimen (J). Bars: 25*µ*m (A, B, D-E’); 10*µ*m (C, F, G, H)

## Discussion

### Primary small undifferentiated neurons form a large neuron population of the early larval brain

Our analysis demonstrates that a large fraction of the neurons of the early *Drosophila* larval brain does not elaborate a branched neurite arbor and synaptic connections. This finding came as a surprise; it had been well established that the large number of neurons produced during the secondary, larval phase of neurogenesis remain undifferentiated until the onset of metamorphosis, resembling in many ways the SU neurons described here (Truman, 1990; Dumstrei et al., 2003; Truman et al., 2004; Pereanu and Hartenstein, 2006), but the same was not assumed for so many of the embryonically generated primary neurons. Previous studies had shown that in the thoracic ganglia, a subset of presumptive adult peptidergic neurons (Veverytsa and Allan, 2012) and motor neurons (Zhou et al., 2009) show a SU phenotype, extending a truncated axon into the peripheral nerve, but failing to form synaptic connections to the musculature. These embryonically born neurons, which transiently express the transcription factor Broad-Z3, differentiate along with the larvally born motor neurons and form dendritic and axonal branches and synapses in the pupa. Due to their delayed differentiation, typical for secondary neurons, Zhou et al. (2009) term these embryonically born thoracic SU neurons “embryonically born secondary neurons”. To avoid confusion, we will stick to the convention that defines all neurons born during the embryonic period as primary neurons, and call them “primary SU neurons”.

The existence of SU neurons (primary or secondary) is most likely tied to the holometabolous life cycle of *Drosophila* where, in terms of structure and function, the larval body (formed in the embryo) differs strongly from the adult body (formed in the larva and pupa). The proliferation of adult-specific cells and organs that takes place in the larva is separated from the differentiated larval structures, possibly in order to prevent interference between novel growth and organ function. In case of the musculature, for example, proliferating adult myoblasts form clusters of cells attached to the peripheral nerves or imaginal discs, outside the larval musculature (Bate, 1993). Neuroblasts generating adult specific neurons are part of the larval brain, but their progeny are arrested in the immature SU state until the onset of metamorphosis, when, under the influence of ecdysone signals, all neurons start branching and generating synaptic connections (Lee et al., 2000; Brown and Truman, 2009; Brown et al., 2006; Veverytsa and Allan, 2012; 2013). If that were not the case, that is, if secondary neurons would continuously differentiate according to their birth date (like regular primary neurons in the embryo), they would constantly and in growing numbers intrude into existing larval circuits, possibly leading to disruptions in functioning of these circuits. It is conceivable that the occurrence of primary SU neurons can be explained by the same reasoning. Based on their (mostly) superficial position, we assume that primary SU neurons are born during the final rounds of embryonic neuroblast divisions. It could be speculated further that there is a “cut-off” line that limits neurons’ ability to commence differentiation, and that this cut-off line falls before the time interval during which primary SU neurons are born, thereby preventing the latter from differentiating.

In support of the notion that the presence of SU neurons is an attribute of holometabolous insects, such cells have not been observed in locusts or other hemitabolans for which neuro-developmental observations have been made. A good number of central neurons of the brain and VNC that were followed throughout development show continuous growth and arborization of their neurite tree (e.g., Bentley and Toroian-Raymond, 1981; Meier et al., 1993). This also includes the unicolumnar neurons of the central complex which, in *Drosophila*, are all born as secondary neurons and undergo a phase of developmental arrest in the larva. In grasshopper, the homologous neurons mature continuously between mid- and late embryonic stages, to form part of a functional central complex right after hatching of the embryo (Boyan and Reichert, 2011).

*Drosophila* SU neurons described in this paper exhibit structural characteristics that are similar to those described for neuronal precursors in the developing vertebrate brain. Thus, postmitotic neuronal precursors of the neocortex or hippocampus, while migrating along radial glia, are small, electron-dense cells with hetero-chromatin-rich nuclei and scant cytoplasm. Typically, they exhibit a bipolar shape, extending a leading and trailing process that are in contact with the radial glia (Eckenhoff and Racic, 1984). The same phenotype is observed in neuronal precursors (“D-cells”) that are generated in the subgranular zone of the hippocampus in adult mammals (Seri et al., 2004; Ngwenya et al., 2006; 2008). As neurons mature, forming dendrites and axons, nuclear and cytoplasmic size increase, and cells become transcriptionally more active, with a concurrent reduction in heterochromatin. Experimental studies have shown that a variety of signaling pathways and receptors for neurotrophic factors become activated by proteins forming part of the complex cell cycle-controlling molecular machinery (Lee et al., 1994; Cicero and Herrup, 2005; Kawauchi et al., 2013). However, the specific mechanism that drives the transition from small, heterochromatin-rich neural precursor to differentiated neuron is little understood. In human and mouse, mutations in the MECP2 protein, which encodes a transcriptional repressor, is associated with a reduction or delay of neuronal maturation (Rett Syndrome; Shahbazian et al., 2002; Martinez de Paz and Ausio, 2017). The gene network (of which MECP2 may form part) that accompanies neural precursor maturation has not been established. In *Drosophila*, this mechanism is embedded into the ecdysone hormonal cycle that controls larval growth and metamorphosis in general. It has been shown that different isoforms of the ecdysone receptor (EcR) are expressed and required for different developmental changes that occur in the nervous system. The EcRB1 and EcRB2 isoforms are expressed in primary neurons that undergo remodeling, including the gamma neurons of the mushroom body (Schubiger et al., 1998; Lee et al., 2000), and blocking this receptor will result in defects of remodeling. In contrast, EcR-A appears to be more dedicated to guide secondary neurons through their maturation and maintenance (reviewed in Brown et al., 2009). The level of ecdysone and its receptors are under the control of developmental paracrine signals, such as in case of the mushroom body cells which produce an activin signal to maintain EcR-B1 levels (Zheng et al., 2003). In addition, intrinsic determinants expressed sequentially in the dividing neuroblasts form part of a feed-back mechanism with the ecdysone cycle (Syed et al., 2017). SU neurons in the *Drosophila* larval brain may present a favorable paradigm to study the process of neuronal maturation downstream of the ecdysone cycle. SU neurons represent a major population at the early larval stage (primary SU neurons) and late larval stage (primary and secondary SU neurons), and can be labeled by specifically expressed factors (e.g., Broad-Z3), which should make them amenable to FACS sorting and systematic gene expression screens.

### SU neurons and the central complex

Most neuropil compartments of the adult *Drosophila* brain have a corresponding larval counterpart (Younossi-Hartenstein et al., 2003; Pereanu et al., 2010); outgrowing fibers of secondary neurons, which form much of the volume of the adult compartments, follow their primary siblings and establish dendritic and axonal branches around this primary scaffold. This principle does not apply to the secondary neurons forming the central complex, for which no anatomically defined larval counterpart exists. Small primordia of the different compartments of the central complex and associated structures (i.e., the bulb and anterior optic tubercle, which relay input to the central complex; Lovick et al., 2017) can be first detected at the late larval stage. These larval primordia of the central complex compartments are formed by the fiber bundles and associated 1lopodia of the secondary lineages which will develop into the central complex of the adult brain. The minute early larval prFB, formed by the FB pioneers described in the present paper, represent an exceptional case. Thus, FB pioneers are primary neurons whose axons extend during the late embryonic phase and gather into a tight commissural bundle located in the center of the crossing fiber masses that constitute the supraesophageal commissure of the early larval brain. Several aspects of the prFB deserve special comment.

(1) From late embryonic stages onward the FB pioneer axons are enclosed by an exclusive glial layer formed by the so called interhemispheric ring glia (Simon et al., 1998). Several pairs of primary glia, located close to the brain midline, make up the interhemispheric ring. Processes of these glial cells assemble into an invariant pattern of sheaths around several individual commissural bundles. Posteriorly, glial processes form two channels, a ventral one containing the great commissure, and a dorsal one, dedicated to the prFB (Lovick et al., 2017; this paper). This dorsal channel conducting the prFB stands out by a pair of glial nuclei attached to its posterior-medial wall (Fig.4); we could unequivocally identify a pair of glial nuclei at that position in the serial EM stack (Fig.4). As the larva grows and secondary tracts of DM1-4 are added to the FB pioneer bundle, the glial channel widens. This volumetric increase continues throughout metamorphosis, and eventually the interhemispheric ring glia accommodates the entire fan-shaped body. Similar to other primary neuropil glia, the interhemispheric ring undergoes apoptotic cell death during mid-pupal stages (Simon et al., 1998; Omoto et al., 2015), and is replaced by a much larger number of small secondary glia that surround the adult fan-shaped body. Genetic studies indicate that interhemispheric glia does play a role in the morphogenesis of the fan-shaped body, even though this role may be relatively minor, or occur late in development. Thus, genetic ablation of glia (Spindler et al., 2009), or loss of function of molecular factors expressed specifically in the interhemispheric ring (Simon et al., 1998; Hitier et al., 2000), result in defective shapes of the adult FB and EB. However, the pathways of DM1-4 lineages at the late larval stage did not display major defects.

(2) FB pioneers differentiate into the pontine neurons of the adult central complex. Pontine neurons differ in their projection from all other columnar neurons, because they connect two columns on either side of the midline. For example, pontine neurons of the right hemispheric DM4 lineage connect the right lateral column of the FB with its left medial column, thereby crossing the midline. In contrast, axons of other, unicolumnar neurons of the right DM4 remain ipsilaterally, connecting only to the right lateral column of the FB (see Fig.1). The trajectories of the FB pioneers re2ect this pontine-typical behavior already in the early larva. Thus, the majority of DM4 SU axons reach the midline and terminate just after crossing it; DM3 axons project slightly further, followed by DM2 and DM1, which reach more than 20*µ*m into the contralateral hemisphere. Outgrowing secondary DM1-4 tracts, even though they initially follow the FB pioneers, show their own characteristic pattern of termination. In particular, secondary axon tracts of DM4 and DM3 do not cross the midline, but form terminal filopodial tufts in the lateral and medial half, respectively, of the ipsilateral prFB (see Fig.5).

(3) Even though primary FB pioneers and their secondary follower tracts are in close contact to each other throughout larval development, ablation of the former does not result in gross structural abnormalities of the latter. Thus, the characteristic trajectories and branching pattern of the Neurotactin-positive secondary tracts of DM1-4 in the late larva lacking FB pioneers appeared indistinguishable from the control. Filopodial tufts of secondary tracts in ablated specimens still assembled into regularly sized globular structures, representing the forerunners of fan-shaped body columns (see Fig.7B). These findings imply that separate guidance systems act on the early born pontine neurons and later born unicolumnar neurons. Nothing is known about the molecular nature of global or local signaling systems controlling the highly ordered architecture of the DM1-4 unicolumnar connections within the central complex. Given that neither ablation of glia, nor loss of primary FB pioneers, causes major changes in this architecture, at least at the initial phase of axonal path1nding, it is likely that local interactions among the different DM1-4 lineages and sublineages plays a predominant role. For example, local repulsion in between neurons of these lineages could be instrumental in specifying the separate, largely non-overlapping medio-lateral domains within the the FB and EB where neurons terminate. Similarly, interactions occurring in between sequentially born sublineages within a given DM lineage could determine the projection to different territories along the anterior-posterior axis. It is not yet known how the different classes of DM neurons distinguished by projection (e.g., PB-FB vs PB-EB vs FB-NO etc) relate to their pedigree, that is, the time they are born, or the sublineage they belong to. It stands to reason that the different intermediate progenitors born from the type II DM neuroblasts are responsible to generate structurally different classes of neurons, but this remains to be con1rmed by detailed clonal analysis. If proven correct, one could surmise that intrinsic factors expressed by a given intermediate progenitor provides its progeny with a specific “projection identity”. Neurons descended from a hypothetical intermediate progenitor A might recognize a more posterior territory within the prFB as their proper destination, whereas neurons formed by a (later born) progenitor B are repelled by the A neurons, and are forced to terminate in more anterior territory. The former class would develop into PB-FB neurons, the later into PB-EB neurons. That repulsive interactions in between sublineages are important has been experimentally proven by a recent analysis of semaphorin signaling in the ellipsoid body (Xie et al., 2017). Here, repulsion among DALv2 R-neurons, born at different times, is instrumental for the proper central>peripheral projection of axons within the EB.

(4) It is an open question what, if any, role the primary neurons of DM1-4 (both differerentiated neurons and other (non-prFB) SU neurons) play in the adult central complex. It is quite possible that these neurons do not contribute at all to this structure; in case of another lineage, DALv2, that has been shown to be the case: secondary DALv2 neurons form the ellipsoid body of the adult brain, but primary DALv2 neurons arborize in the lateral accessory complex (LAL) and inferior protocerebrum (IPa) of the larva and adult, but do not become part of the ellipsoid body (Lovick et al., 2017). The same may be true for the primary neurons of DM1-4. On the other hand, at least a (small) subset of DM4 definitely will be incorporated into the central complex: the dopaminergic neurons of the PPM3 cluster, which profusely innervate the central complex and its associated structures (LAL, BU) have been identified as primary neurons of DM4 (Hartenstein et al., 2017). The arborization pattern and connectivity of primary DM1-4 neurons (as that of primary neurons in general) will be worked out in the near future, based on the same serial EM stack that served as the basis for the current work; however, additional markers that remain continuously expressed in primary neurons from larval to adult stages will help solving the puzzle of how these neurons are reorganized during metamorphosis and what fate awaits these neurons in the adult brain.

Aside from pioneering the fan-shaped body primordium, SU neurons form part of almost all lineages of the early larval brain, but we do not yet know what fate awaits these neurons. In view of the case represented by the FB pioneers, we surmise that other SU neurons also survive and differentiate during metamorphosis. The axonal projection of SU neurons of a given lineage pre1gures (generally in a rudimentary way) the pathway formed by later born secondary neurons of that lineage (see Figure 3). Markers similar to the one provided by R45F08-Gal4 are required to establish what type of adult neurons the different SU neurons give rise to. Of particular interest are SU neurons of lineages that, like DM1-4, contribute to the adult central complex. In case of DALv2, which generates all of the R-neurons of the EB, a single SU neuron exists in each hemisphere (see Table 1). This neuron projects a short axon along the primary tract, but does not reach the EB primordium described for the late larval stage in previous works (Lovick et al., 2016; 2017). Two other lineages, DALcl1 and DALcl2, contribute a large number of secondary neurons to the anterior visual pathway, which provides input to the central complex (Omoto et al., 2017). Both lineages are composed of two hemilineages, DALcl1/2d (dorsal) and DALcl1/2v (ventral). DALcl1/2d differentiate into small neurons whose proximal dendrites innervate the anterior optic tubercle, and distal axons the bulb, where they target the dendrites of DALv2 neurons (Omoto et al., 2017; Lovick et al., 2017). DALcl1/2 form a relatively large number of SU neurons which for the most part follow the dorsal pathway, suggesting that they belong to the DALcl1/2d hemilineage. As outlined above, secondary neurons of DALcl1/2 innervate the anterior optic tubercle and the bulb of the adult brain. Do the earlier born primary SU neurons form early larval primordia of these compartments, analogous to the prFB established by SU neurons of DM1-4. Most DALcl1/2 SU neurons extend relatively short axons that follow the differentiated DALcl1/2d neurons, cross the peduncle of the mushroom body, and terminate shortly thereafter. A few other DALcl1/2 SU neurons project further, but terminate at different locations along the primary tract. In other words, a spatially restricted territory that houses all DALcl1/2d SU terminations, and that might therefore be considered a forerunner of the bulb, does not exist in the early larva. Similarly, no projections of SU neurons are concentrated in a region that might correspond to the primordium of the anterior optic tubercle. In conclusion, SU neurons of lineages DM1-4 may represent a rare case where primary neurons establish a blueprint for an adult-specific brain compartment.

## Methods and Materials

### *Drosophila* Stocks

The following *Drosophila* lines were acquired from the Bloomington *Drosophila* Stock Center (BDSC), followed by citation and stock number in parentheses when applicable: R45F08-Gal4 (Jenett et al., 2012; *#*49565), 10xUAS-mCD8::GFP (Pfeiffer et al., 2010; *#*32186), R57C10-Flp-2::PEST,su(Hw)attP8::HA_V5_FLA (Nern et al., 2015; *#*64089), UAS-hid UAS-rpr UAS-lacZ (Zhou et al.,1997), Sco/CyO; tub-Gal80ts (Bloomington *#*7018). Drl-tau::lacZ (Chen and Hing, 2008), was kindly provided by H. Hing, The College at Brockport, State University of New York, New York.

### Selective ablation of fan-shaped body pioneers

FB pioneers were ablated from late first instar to second instar. The ablation experiment was carried out using the following genotype: UAS-hid UAS-rpr UAS-lacZ/+;10xUAS-mCD8::GFP/+; R45F08-Gal4/tub-Gal80ts. Freshly hatched larvae were collected and kept at 18°C degrees for two days (corresponding to 24 hours at 25 °C). Subsequently, these late first instar/early second instar larvae were transferred to 29°C for 24 hours. After the temperature shift, larvae were transferred back to 18 °C. When reaching the third instar wandering stage, late larvae were collected, dissected and processed for analysis. The constitutive activation of UAS-hid,rpr by R45F08-Gal4 is lethal, and only survives into larval stages with Gal80-mediated suppression. Control flies were either males lacking UAS-hid UAS-rpr UAS-lacZ and tub-Gal80ts (Fig.7A-C), or females lacking a temperature shift and maintained at 18 degrees, which were completely viable (data not shown). Most experimental female larvae (heat-shifted) exhibited lethality; a small fraction were slowed in their development relative to male controls, and this was the cohort utilized for analysis.

### Single cell clones

Single-cell analysis of the differentiated SU neurons in the adult was conducted using the multicolor 2ip-out (MCFO) method described previously (Nern et al., 2015). Flies of the genotype R57C10-Flp-2::PEST,su(Hw)attP8::HA_V5_FLAG_1 were crossed to R45F08-Gal4 to generate the required progeny. The cross was kept at 25°C under normal light/dark cycle. 1-3 day adults were dissected, stained and imaged to screen for single-cell labeling of R45F08-Gal4-positive pontine neurons.

### Immunohistochemistry

Antibodies: The following primary antibodies were obtained from Developmental Studies Hybridoma Bank, DHSB, (Iowa City, IA): rat anti-DN-cadherin (DN-EX *#*8, adult, third instar larva: 1:8, first instar larva: 1:10), mouse anti-Brp (nc82,1:20), mouse anti-Neurotactin (BP106, 1: 6) mouse anti-Neuroglian (BP104, Adult: 1:12, first instar larva: 1:14), mouse anti-repo (1:10), rabbit anti-*β*Gal (1:3). We also used rabbit anti-HA (1:300, Cell Signaling Technologies), chicken anti-GFP (1:1000, Abcam) and rat anti-FLAG (1:300, Novus Biologicals). The following secondary antibodies were obtained from Jackson ImmunoResearch; Molecular Probes) and used: Alexa 568-conjugated anti-mouse (1:500), cy5-conjugated anti-mouse (1:300), cy3-conjugated anti-rat (1:300), Alexa 488-conjugated anti-rabbit (1:1000), cy5-conjugated anti-rat (1:300), cy3-conjugated anti-rabbit (1:160), and Alexa 488-conjugated anti-rabbit (1:300). We also obtained Alexa 488-conjugated anti-chicken (1:1000) from Thermo Fisher Scientific.

Adult brain preparation: Brain samples were dissected in phosphate buffered saline (1X PBS, pH 7.4) for 30 minutes. The dissected brains were then fixed in ice-cold 4% formaldehyde fixing solution [750*µ*l phosphate buffered saline (PBS); 250*µ*l 16% paraformaldehyde, EM grade (Sigma)] for 2.5 hours before washing four times, 15 minutes each, with ice-cold 1X PBS. Then the brains were dehydrated in a series of ice-cold ethanol diluted with 1X PBS (5,10,20, 50, 70 and 100% ethanol; for 5 minutes at each concentration). After hydrating, the brains were stored in 100% ethanol at -20 °C overnight. For rehydration, we repeated the hydration process in reverse order. These rehydrated brains then were washed with ice-cold 1X PBS 4 times, 15 minutes each. Next, they were washed 3 times, 15 minutes each, with ice-cold 0.3% PBT (PBS; 0.3% Triton X-100, pH7.4), followed by 4 washes, 15 minutes each, on the nutator at room temperature. After washing samples were blocked (10% normal goat serum in 0.3% PBT) and then stained with primary and secondary antibodies for 3 days each under gentle shaking on the nutator at 4 °C. Washes were done when replacing primary with secondary antibodies with 0.3% PBT, and when removing secondary antibodies. Finally samples were transferred to Vectashield, mounted and imaged. Brains with MCFO single cell clones were processed following a shortened protocol in which the dehydration/rehydration steps were omitted.

Larval brain preparation: Larval brains were dissected at the indicated age (24, 48, 60, 72 and 96 hours old) in phosphate buffered saline (1X PBS, pH 7.4) for 30 minutes and then fixed in 4% formaldehyde 1xing solution (900 *µ*l 1X PBS, 100 *µ*l 37% paraformaldehyde) for 30 minutes with gentle shaking. The fixed brains then were washed with the indicated buffer solution 4 times, 15 minutes each. Brains were also subjected to the methanol dehydration series diluted with 1X PBS (10 minutes in 25% and 50% methanol each, and 4x 1-minute washes in 75% methanol, and one 5-minute wash in 100% methanol). Samples were then stored in 100% methanol at -20 °C overnight, then rehydrated the next day by washing with 1X PBS once. Subsequently samples were washed with 0.1% PBT (PBS 0.1% Triton X-100, pH7.4) on the nutator. After washing, samples were blocked (10% normal goat serum in 0.1% PBT for 30 minutes), and incubated in primary and secondary antibodies for 2 days each at 4°C while gently shaking on the nutator. Washes with 0.1% PBT were done when replacing primary with secondary antibody and before placing the samples in VectaShield.

### Digital reconstruction and analysis of SU neurons from serial EM

SU neurons were reconstructed from a series of TEM sections of an entire nervous system of a 6 hour old larva as described in Ohyama et al. (2015). Sections were imaged semi-automatically using the Leginon software (Suloway et al., 2005) and assembled using the TrakEM2 (Cardona et al., 2012). Reconstruction utilized CATMAID, a web-based software developed (Saalfeld et al., 2009) with updated neuronal skeletization and analytic tools (Schneider-Mizell et al., 2016).Reconstructed using the method described in Ohyama et al., (2015); Schneider-Mizell et al. (2016). For obtaining the area of the nucleus and cytoplasm of a cell, a section at or near the center of the soma was chosen. Subsequently, the length of the longest and shortest radius was obtained by using a tool of CATMAID this gives the radius in nm. The average radius was taken to calculate area.

## Acknowledgments

We thank Akira Fushiki, Ivan Larderet, Feng Li, Avinash Khandelwal, Timo Saumweber, Jennifer Lovick, Larisa Meier, Katharina Eichler, Javier Valdez-Aleman, Simon Sprecher and Philipp Schlegel for their contributions in the reconstruction 17.2% of SU neurons.

## Supplementary Figures

### Supplementary Figure 1

**Fig.S1.**
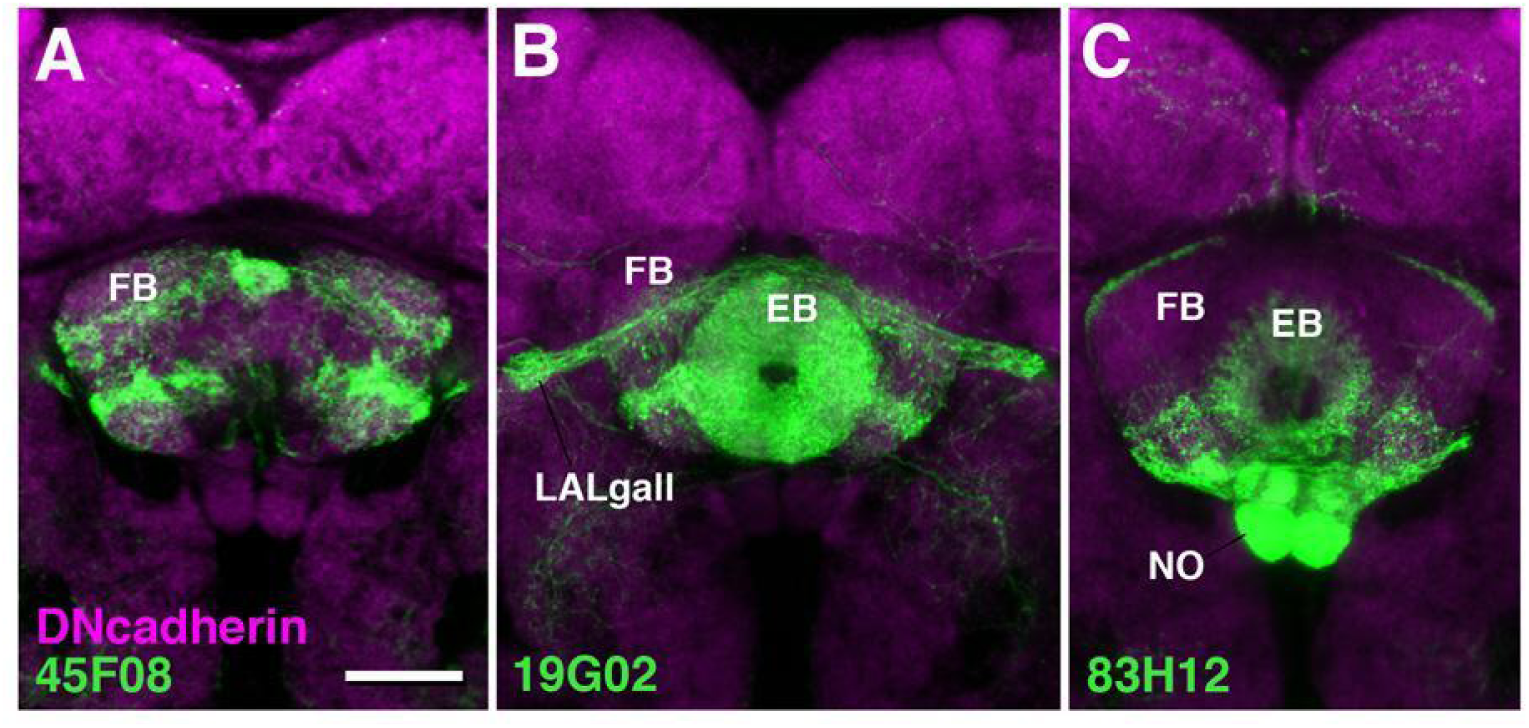
Z-projection of frontal confocal sections of median region of adult brain, showing projection of driver lines R45F08-Gal4 (pontine neurons; A), R19G02-Gal4 (PB-FB-LALgall; PB-EB-LALgall), and R83H12-Gal4 (PB-FB-NO) in fan-shaped body (FB), ellipsoid body (EB), and noduli (NO). Bar: 20*µ*m

### Supplementary Figures S2-S9

Graphic representation of all SU neurons of the first instar (L1) larval brain ordered by lineage groups. SU neurons of a given lineage are clustered in a cohesive tract, the primary axon tract. These tracts provide useful landmarks for the ongoing analyses of the L1 brain circuitry using the CATMAID L1 serial TEM dataset, which is the rationale for preparing this set of supplementary 1gures.

Each of the figures is structured similarly. Panels (A-C) show 3D digital models of L1 brain in dorsal view (A; anterior to the top), lateral view (B; anterior to the left, dorsal up) and anterior or posterior view (C; dorsal up). The neuropil surface is rendered in light grey; mushroom body, located in center of brain neuropil, is outlined in dark grey. In Figures S2–S4 and S7-S9 two additional landmarks, the antennal lobe (antero-ventral brain neuropil) and larval optic neuropil (postero-lateral brain neuropil), are represented by renderings of individual neurons [larval optic neuropil: GlulOLP (Larderet et al 2017); antennal lobe: Broad D1 (Berck et al 2017)]. Cell bodies and axons of all SU neurons of both hemispheres are shown as colored spheres and lines, respectively. Differential coloring indicates association of SU neurons with discrete lineages, as identified in color key presented in the table at the bottom right of each figure. Small annotated circles indicate location where corresponding primary tracts enter the neuropil. Annotations indicate the lineage name. In many cases, only the last part of the lineage signi1er is shown for clarity sake; for example, in Fig.S2, the pair “BAla1/2” is abbreviated as “la1/2”. The x,y,z spatial coordinates of entry points shown in panels A-C are given in table at bottom right of each 1gure. The origin (z=0/x=0/y=0) represents the anterior/right/dorsal corner of the virtual cuboid that encloses the data set. The units for z is section number (z=0 corresponding to most anterior section; section thickness = 60nm); x and y indicate pixels, with each pixel corresponding to approximately 3.5nm. For each lineage, upper row of numbers present coordinates of tract entry point in right hemisphere (R); lower row gives coordinates for left entry point. For nomenclature and grouping of lineages, see Lovick et al. (2013) and Hartenstein et al. (2016).

Abbreviations: AL antennal lobe; CX calyx; ML medial lobe; PED peduncle; VL vertical lobe; VNC ventral nerve cord

**Fig.S2.**
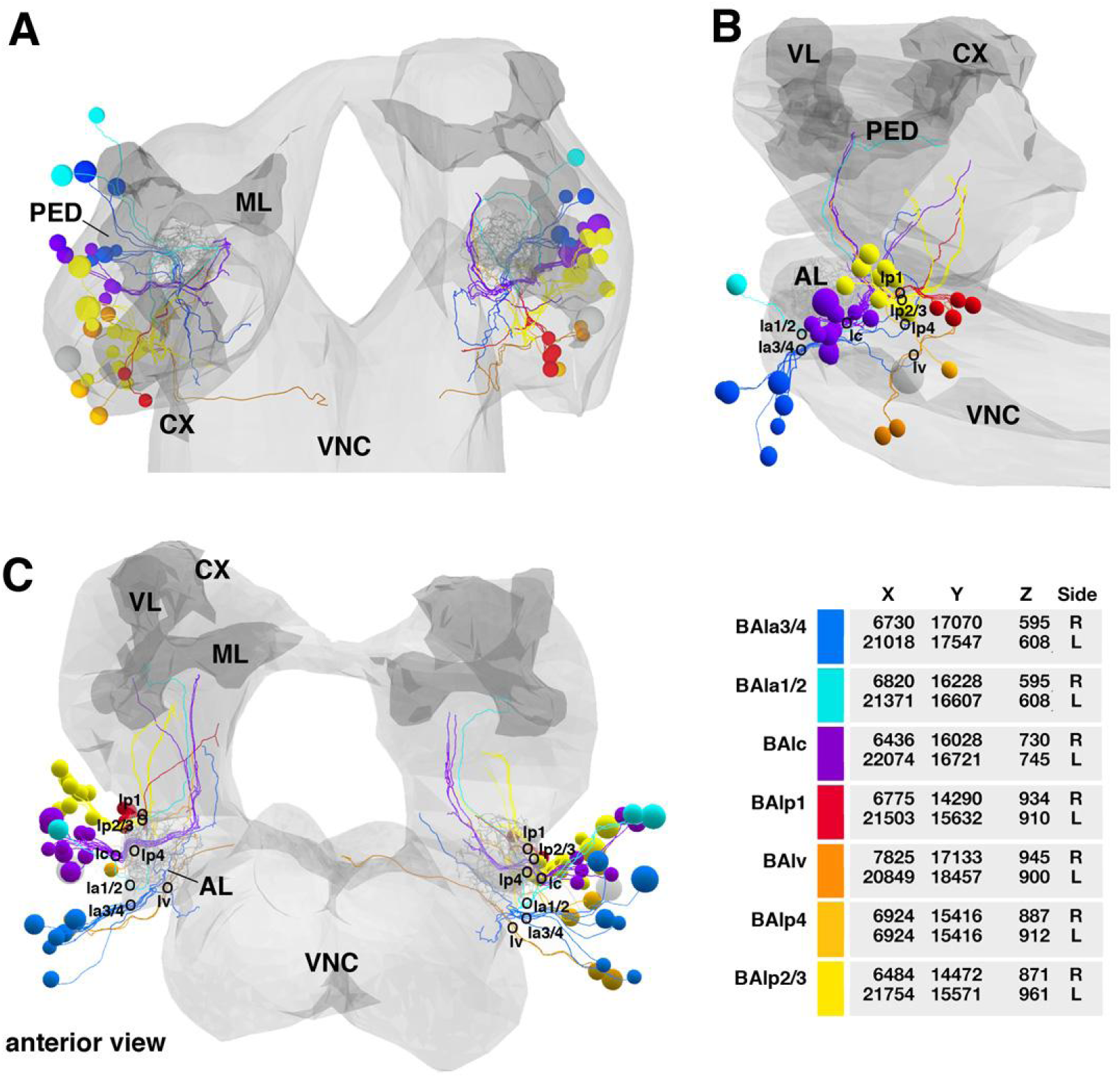
Lateral baso-anterior group of lineages (BA)

**Fig.S3.**
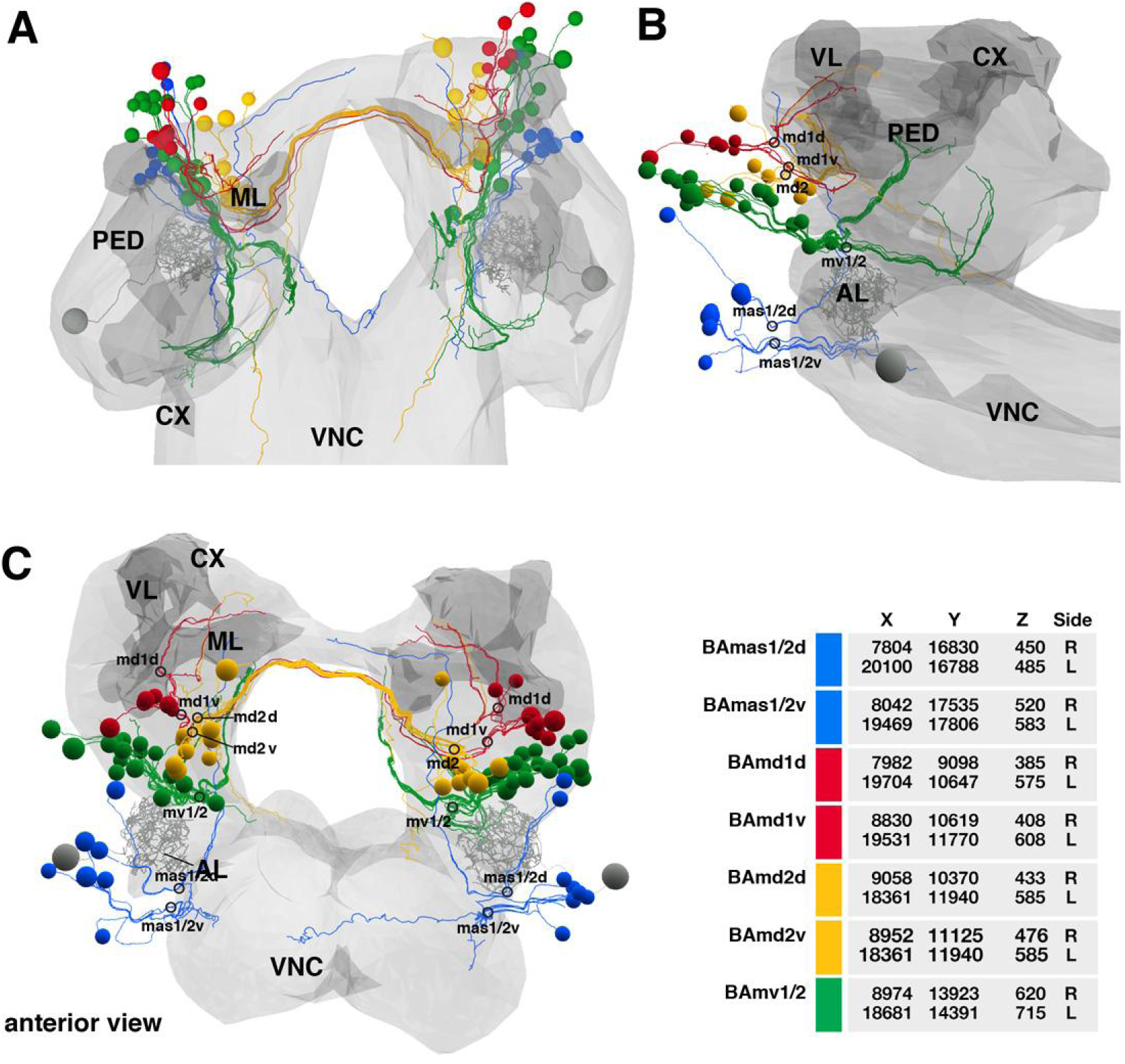
Medial baso-anterior group of lineages (BA)

**Fig.S4.**
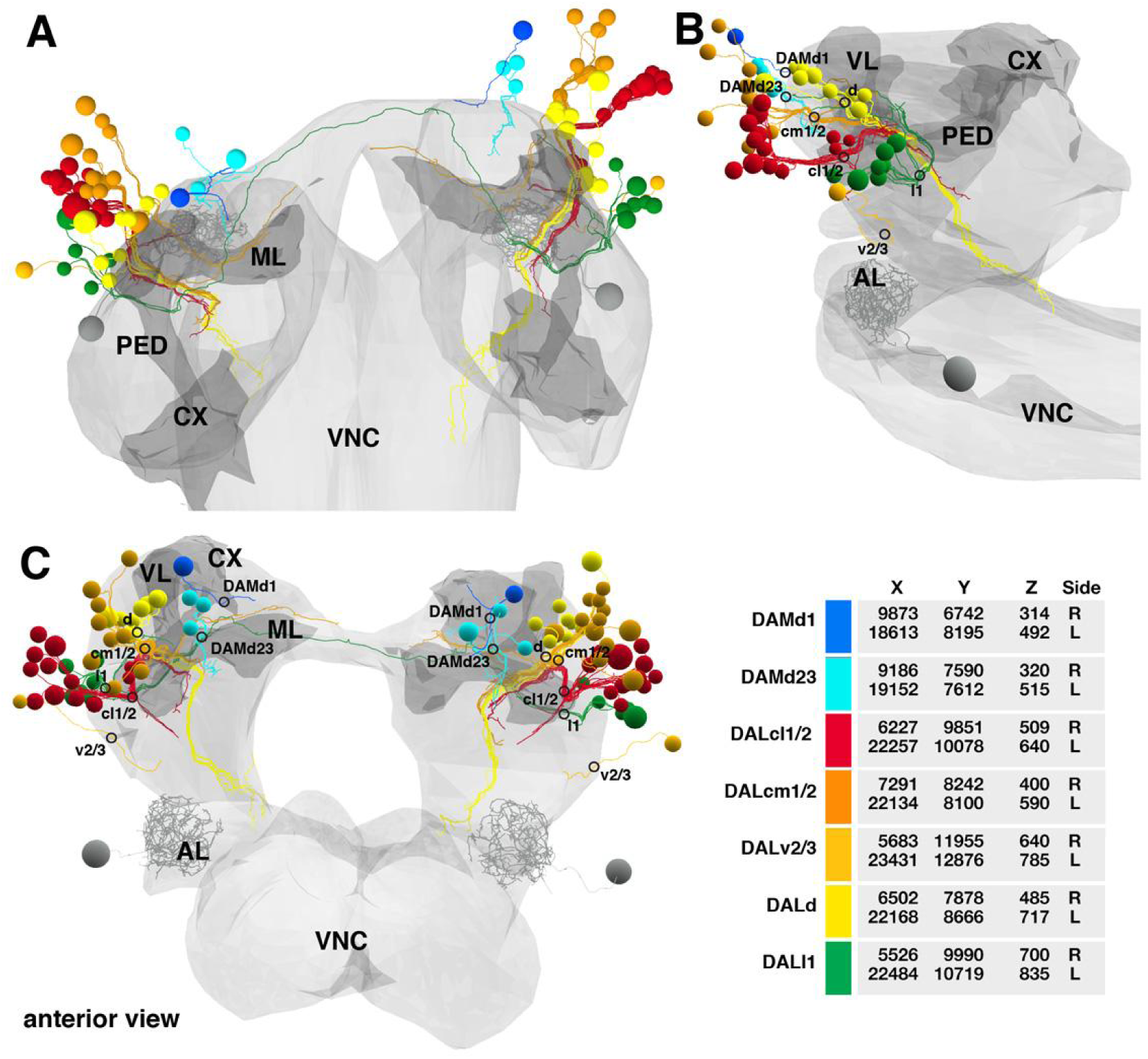
Lateral dorso-anterior and medial dorso-anterior groups of lineages (DAL, DAM)

**Fig.S5.**
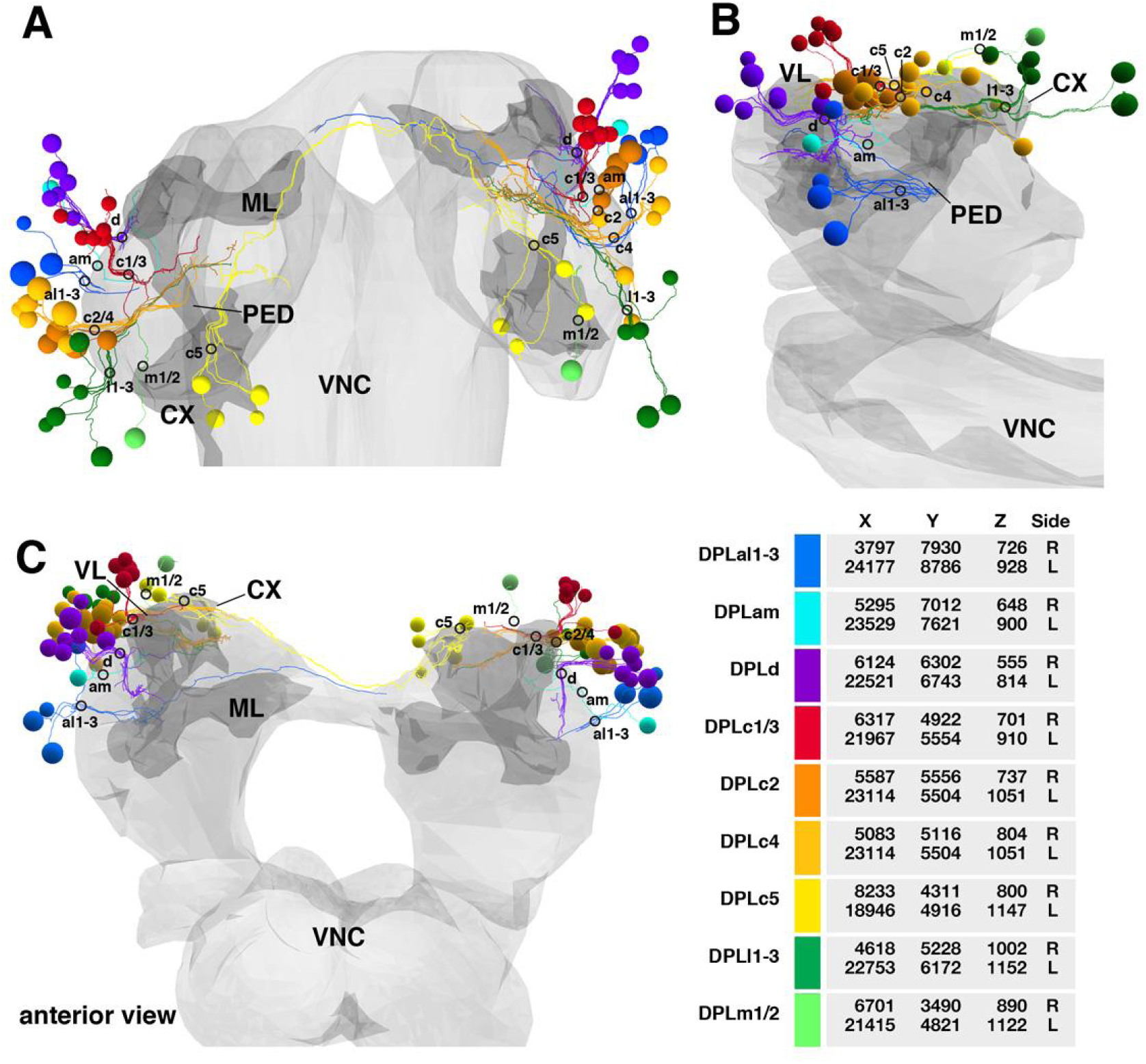
Lateral dorso-posterior group of lineages (DPL)

**Fig.S6.**
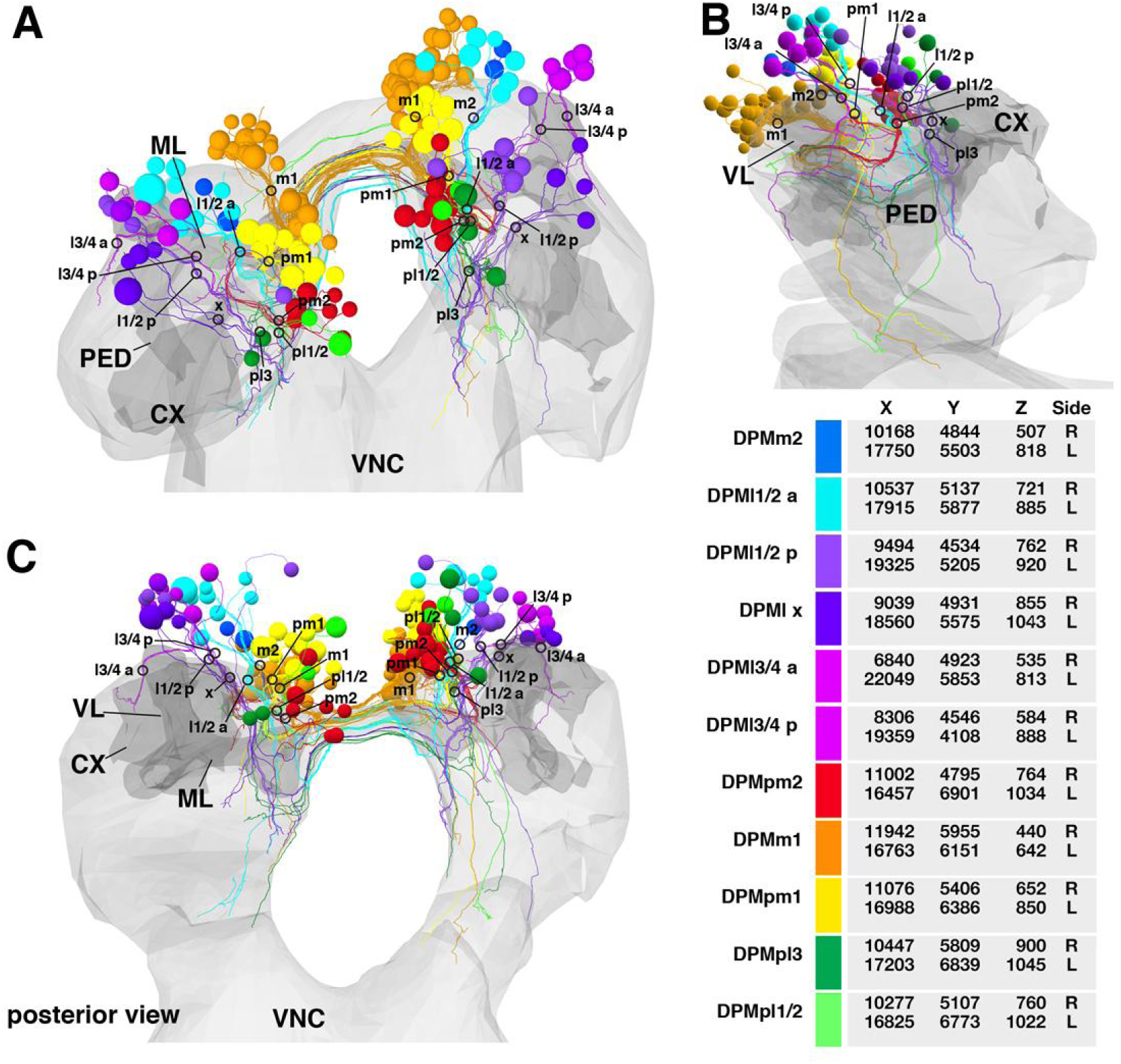
Medial dorso-posterior group of lineages (DPM)

**Fig.S7.**
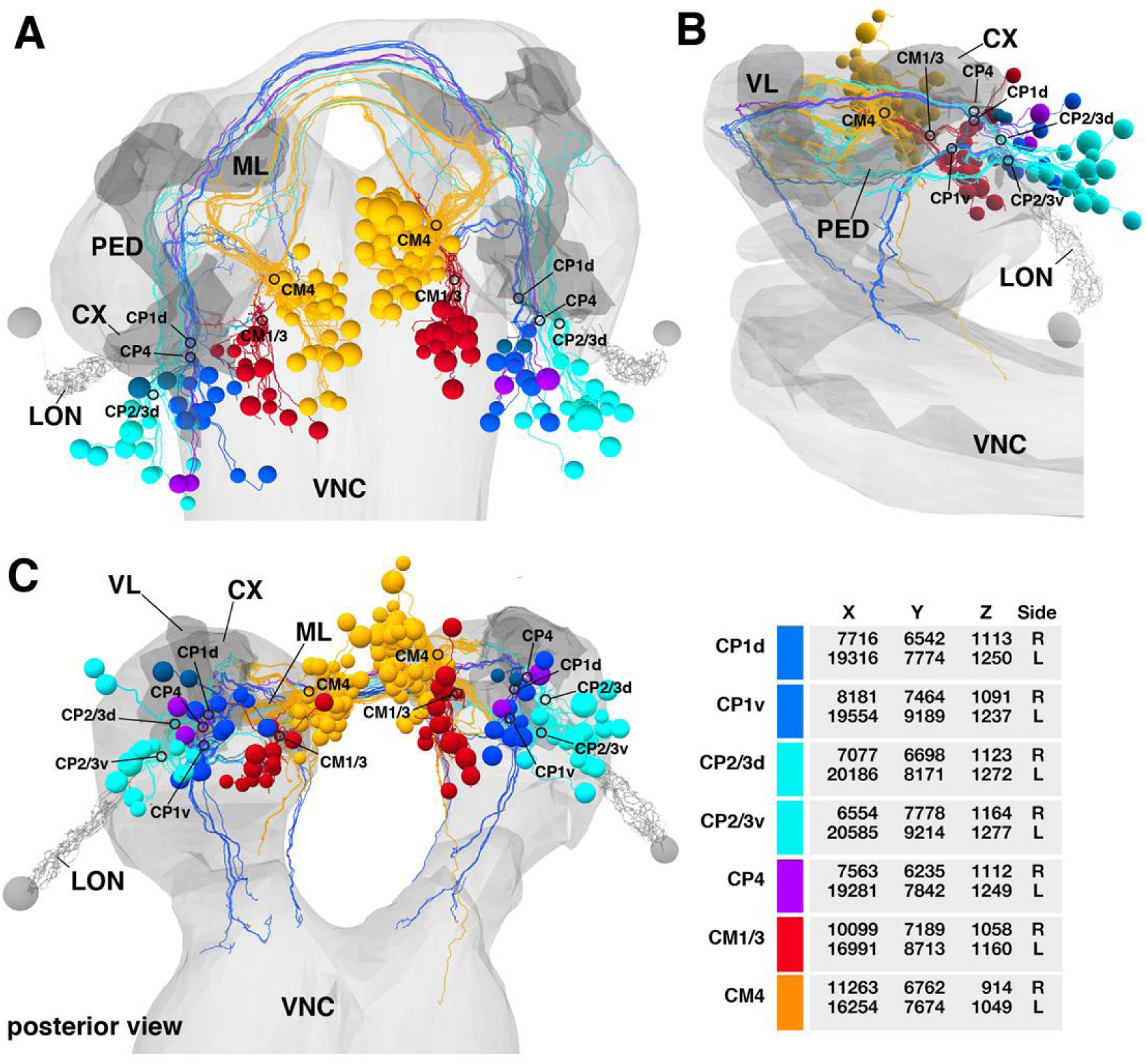
Lateral centro-posterior and medial centro-posterior groups of lineages (CP, CM)

**Fig.S8.**
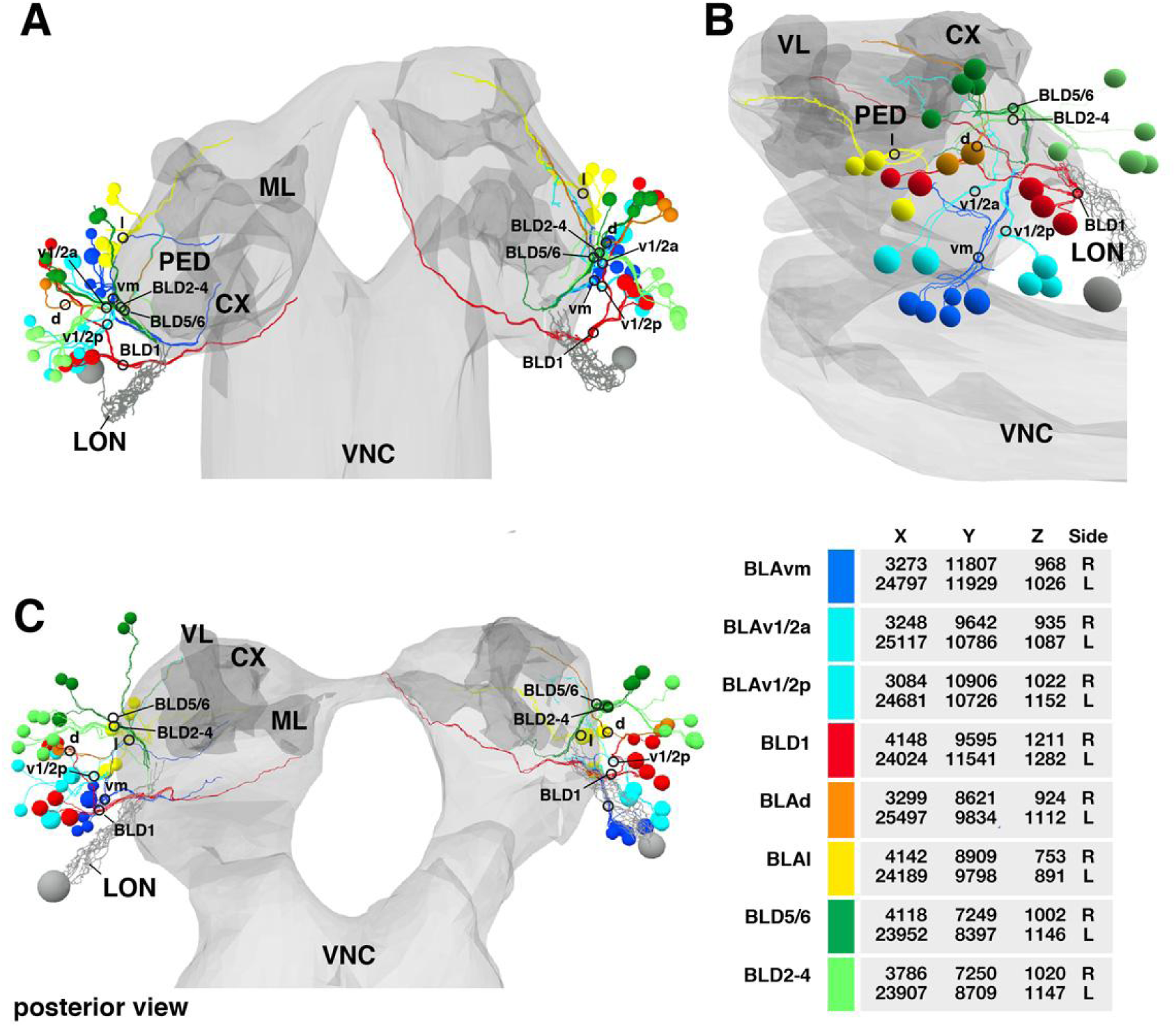
Anterior baso-lateral and dorsal baso-lateral groups of lineages (BLA, BLD)

**Fig.S9.**
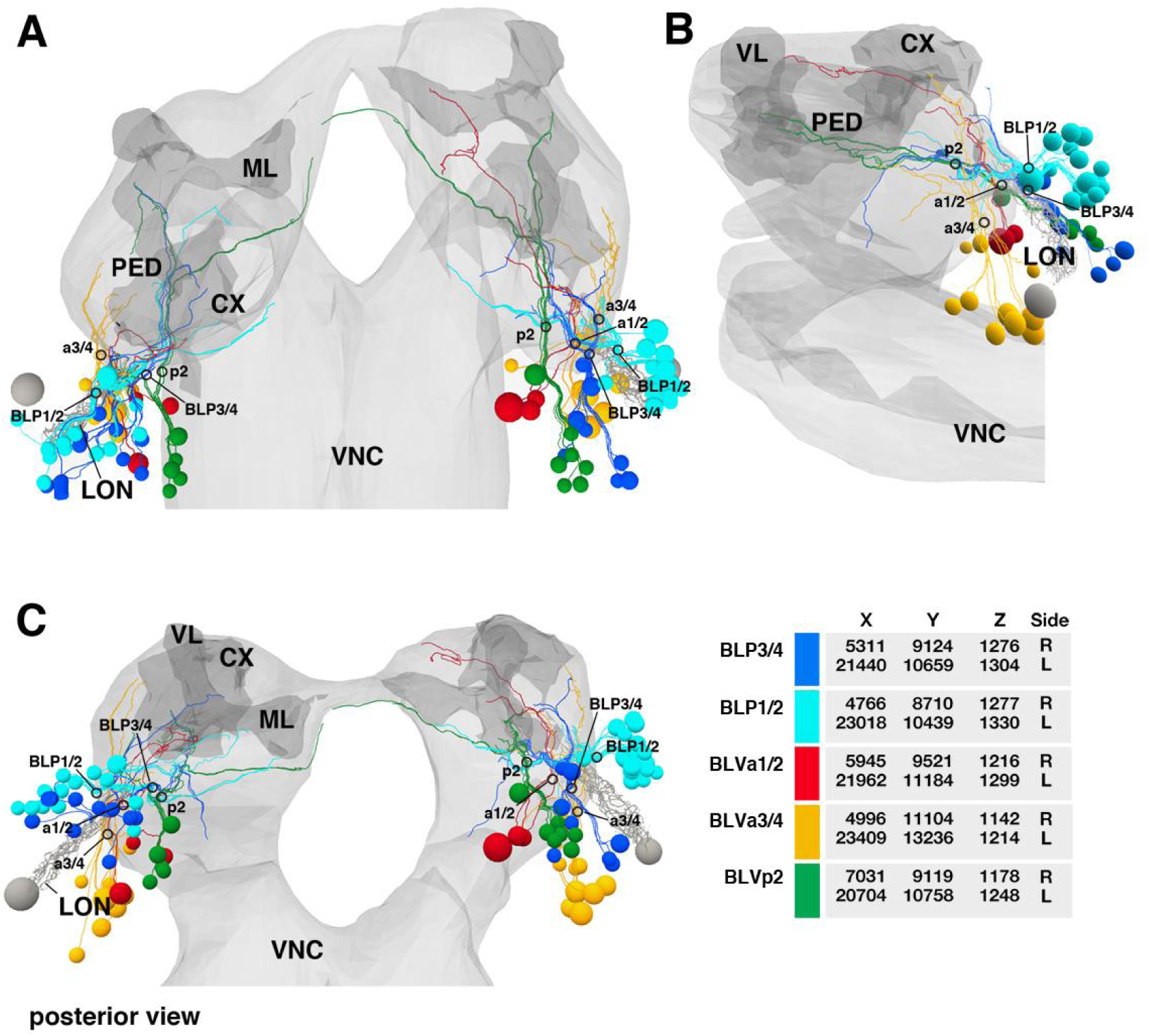
Posterior baso-lateral and ventral baso-lateral groups of lineages (BLP, BLV)

